# An in-cell approach to evaluate E3 ligases for use in targeted protein degradation

**DOI:** 10.1101/2024.12.19.629291

**Authors:** Yunan Zheng, Anamika Singh, Zeqi Niu, Violeta Marin, Jonathon Young, Paul Richardson, Marcus L. Hemshorn, Richard B. Cooley, P. Andrew Karplus, Scott Warder, Anil Vasudevan, Justin M. Reitsma, Ryan A. Mehl

## Abstract

One of the major challenges in evaluating the suitability of potential ∼700 E3 ligases for target protein degradation (TPD) is the lack of binders specific to each E3 ligase. Here we apply genetic code expansion (GCE) to encode a tetrazine-containing non-canonical amino acid (Tet-ncAA) site-specifically into the E3 ligase, which can be conjugated with strained trans-cyclooctene (sTCO) tethered to a neo-substrate protein binder by click chemistry within living cells. The resulting E3 ligase minimally modified and functionalized in an E3-ligand free (ELF) manner, can be evaluated for TPD of the neo-substrate. We demonstrate that CRBN encoded with clickable Tet-ncAA, either in the known immunomodulatory drug (IMiD)-binding pocket or across surface, can be covalently tethered to sTCO-linker-JQ1 and recruit BRD2/4 for CRBN mediated degradation, indicating the high plasticity of CRBN for TPD. The degradation efficiency is dependent on location of the Tet-ncAA encoding on CRBN as well as the length of the linker, showing the capability of this approach to map the surface of E3 ligase for identifying optimal TPD pockets. This ELF-degrader approach has the advantages of not only maintaining the native state of E3 ligase, but also allowing the interrogation of E3 ligases and target protein partners under intracellular conditions and can be applied to any known E3 ligase.

## Introduction

A healthy cell maintains an exquisite balance of the synthesis and degradation rates of its various proteins via a set of complex processes that together are known as proteostasis (i.e. protein homeostasis). This governing of the integrity of the cellular proteome involves controlling multiple interconnected pathways responsible for protein synthesis, folding, transport, and disposal^1, 2, 3^. And many diseases, including cancer, neurological conditions, metabolic diseases, and age-related pathologies have been linked to dysregulated proteostasis ^4, 5, 6^.

In eukaryotes, one key process of proteostasis is the ubiquitin-proteasome system (UPS), in which added ubiquitin tags targets specific damaged, misfolded, or no longer needed proteins for proteasomal degradation ^7^. The discovery of the UPS was recognized by the 2004 Nobel prize in chemistry ^8^, and numerous studies have highlighted its importance for cellular health ^9, 10, 11^, and ^12^). The UPS-mediated protein degradation pathway requires the coordination of ubiquitin-activating enzymes (E1), ubiquitin-conjugating enzymes (E2), and ubiquitin-protein ligases (E3). E3 ligases recognize targets through degrons and facilitate the transfer of ubiquitin. Based on their mechanisms of ubiquitin transfer and domain architecture, E3 ligases can be classified into three main families ^13^: the RING (really interesting new gene) finger family, the HECT (homologous to E6AP C-terminus) family, and the ―RING-between-RING‖ (RBR) family. The RING finger subfamilies, Cullin-RING ligases (CRLs), make up over 40% of all E3 ligases. CRLs are modular enzymes that consist of four components: a scaffold cullin protein, a RING finger protein for engaging an E2 enzyme, a substrate receptor for target recognition, and adaptor proteins linking the receptor to cullin^14, 15, 16^. The most widely recognized CRL is cereblon (CRBN), which functions as the substrate receptor for the CRL4 complex (Fig. 1a).

**Fig 1:**
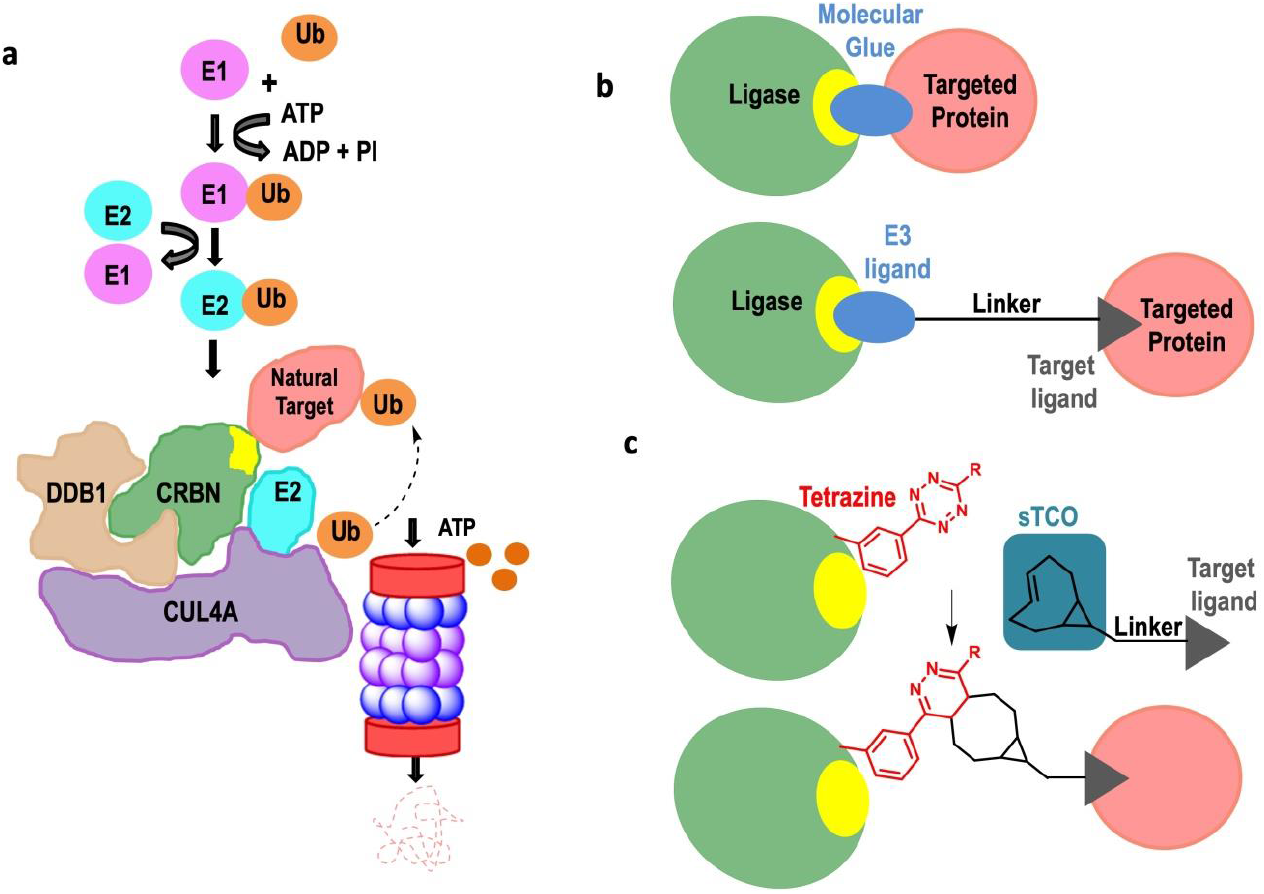
Strategies for targeted protein degradation. (a) Steps in the natural Ubiquitin Protease System (UPS) highlighting the role of the E3 ligase (green) in recognizing the target protein (pink) that is destined for ubiquitination and degradation. (b) How a molecular glue (top) and bi-functional PROTAC molecule (bottom) bring an E3 ligase and a neosubstrate target protein into proximity and triggering its ubiquitination and degradation. (c) Concept for how an E3-ligand free (ELF) degrader (consisting of a trans-cyclooctene group linked to a target protein ligand), after a click chemistry reaction with an E3 ligase containing a tetrazine moiety, can mimic the functionality of a PROTAC molecule in bringing an E3 ligase and target protein together.

Over the past two decades, effective therapeutics have been developed that coopt the UPS to degrade specific disease-causing proteins ^17, 18^, and these powerful targeted protein degradation (TPD) strategies often target otherwise ―undruggable‖ proteins ^19, 20, 21^. One approach involves small molecule ―molecular glue‖ compounds ^22, 23, 24^ which bind tightly to an E3 ligase, changing its surface in way that creates a binding site for another protein which when bound gets ubiquitinated and targeted for degradation (Fig. 1b). Classic examples are immunomodulatory drugs (IMiDs) like thalidomide that ―glue‖ the E3 ligase CRBN to certain transcription factors ^25, 26, 27^ and cause their proteasome-dependent degradation ^19, 26, 28, 29^. A more versatile TPD approach using proteolysis-targeting chimeras (PROTACs) was developed by Sakamoto *et al*. ^30^. PROTAC molecules consist of a linker connecting a ligand for a specific E3 ligase with a ligand for a target protein of choice, which brings the target proteins close to the E3 ligase (Fig. 1b) and leads to the ubiquitination of the target protein ^31, 32^.

Thus far TPD applications have been limited to using just a dozen of the ∼700 human E3 ligases ^33, 34, 35^, because there are only a few for which specific ligand binding sites and high-affinity ligands have been identified ^36, 37^. The main ―workhorses‖ are CRBN and the von Hippel-Lindau (VHL) E3 ligases; these are both well characterized, have known tight-binding ligands, and have been successfully redirected to degrade hundreds of proteins ^20, 38^. Despite these therapeutic advances, we still know very little about the structural constraints required for E3-guided protein degradation including which potential surface locations on E3 ligases are amenable for developing molecular glue type therapies.

The current state-of-the-art to validate E3 ligases for TPD relies on either 1) engineering the ligase with protein tags to allow tag-induced proximity that brings E3 ligase and target protein in proximity upon strong interaction of the protein tags (e.g., HaloTag/FKBP12, dTAG, aTAG or GFP nanobody) ^39, 40, 41^ or 2) producing a recombinant E3 ligase and covalently functionalizing it *in vitro* with a neo-substrate protein binder based on cysteine-maleimide *<u>co</u>*valent modification *f*ollowed by *<u>E</u>*3 *<u>e</u>*lectroporation (COFFEE) back into cells ^36, 42^. The limitations of protein tag-induced proximity are that site-specificity cannot be control and the bulky 15∼30kDa protein tags that are restricted to N or C termini attachment. While COFFEE involves a less-perturbing modification of E3 ligases, only those that can be purified, labeled efficiently on a single cysteine and returned into functional cellular UPS complexes can be explored. Due to these limitations, neither approach can shed light on the structural plasticity and potential ligand binding sites of the untapped E3 ligases.

Since genetic code expansion (GCE) ^43^ using non-canonical amino acids (ncAAs) can preserve intracellular ligase function and allow for residue-specific introduction of click-chemistry ligation sites, GCE presents an avenue to overcome the above limitations. Indeed, by exploiting the tetrazine (Tet) – transcyclooctene (TCO) click-chemistry reaction which can give rapid, quantitative ligation within live eukaryotic cells ^44^, it should be possible to position a Tet ncAA anywhere on the surface of an E3 protein, allowing one to survey the surface and map areas that are optimal for generating protein complexes that lead to degradation. If proven correct, this approach could be applied universally to all E3 ligases for exploring not only their potential for neo-substrate TPD, but also their ―hot pockets‖ to gain confidence for chemistry campaign to generate a specific binder. In this approach, the live cells containing E3-Tet proteins would initiate target protein degradation when exposed to a cyclopropene-fused transcyclooctene (sTCO) reagent attached to a variable linker and a ligand specific for a target protein (Fig. 1c). We call this an E3-ligand free-Degrader or ELF-Degrader approach.

Here, we test this ELF-Degrader approach in a human cell line by evaluating the TPD efficiencies of multiple Tet surface positions of the E3-ligase CRBN reacted with an sTCO-fused JQ1, a ligand used in the successful PROTAC work targeting the neo-substrate BRD2/4 ^45, 46^. We discover a remarkably high level of plasticity with regard to the surfaces of CRBN that can direct efficient protein degradation using ELF-Degraders, with longer linkers (within the range tested) generally performing better.

## RESULTS

### Strategy for development of a GCE-based E3-ligand free TPD

As a model system for developing the ELF-Degrader approach to TPD (Fig. 1c), we sought to recapitulate the efficient CRBN-mediated ubiquitination and proteasomal degradation of the bromodomains BRD2 and BRD4 (BRD2/4) that is induced by the dBET6 PROTAC molecule – a chimera with the BRD2/4 ligand JQ1 conjugated to the CRBN binder pomalidomide ^47^. For the E3 anchor chemistry, we chose the tetrazine-3-butyl non-canonical amino acid (Tet3Bu; Fig. 2a) since it incorporates well in HEK293T cells, undergoes rapid in-cell click reactions with cell permeable sTCO reagents ^44^, and its on-protein reactivity can be readily assessed by the fraction of protein undergoing a gel mobility shift after an *in vitro* reaction with an sTCO-PEG_5K_ polymer ^48, 49^

**Fig 2:**
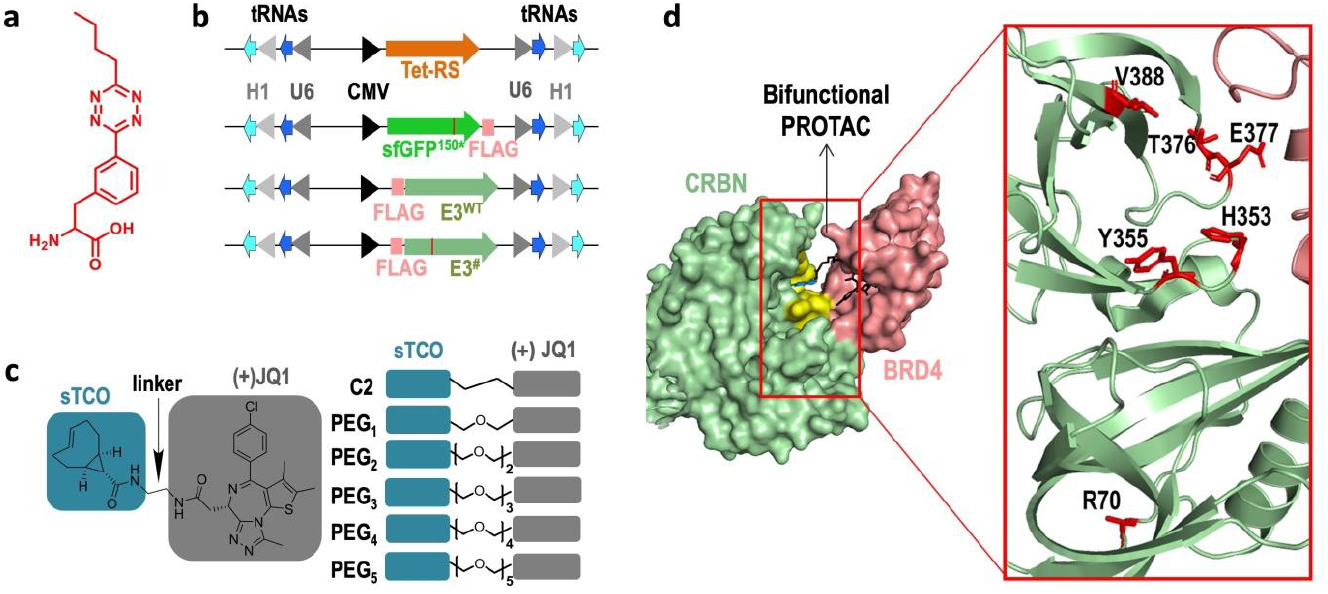
Components used in developing the ELF degrader approach. (a) Chemical structure of the tetrazine3.0-butyl (Tet3Bu) ncAA, shown in red throughout this paper. (b) Plasmid constructs with GCE machinery (NES-RS and tRNAs), control sfGFP150, and protein of interest (CRBN) genes with FLAG-tags as shown and promoters. Also noted are cytomegalovirus (CMV), and H1 and U6 tRNA promoters. (c) Chimeric ELF degrader reagents all have strained trans-cyclooctene (blue box) at one end and a JQ1 target protein ligand (grey box) at the other. Right-hand part names and shows the seven linkers used. (d) Space-filling image of the complex between Cereblon (green with yellow iMiD binding pocket), BRD4 (pink) and the bifunctional PROTAC dBET6 (black sticks), (PDB code: 6BN7)^47^. Close up of interface shows ribbon diagram highlighting the six CRBN residues (red stick models) initially selected for tetrazine incorporation.

The GCE system for efficiently expressing Tet3Bu-containing proteins in HEK293T cells ^44^ includes two plasmids, one expressing the Tet3Bu-specific aaRS, the other expressing the target protein for Tet3Bu incorporation, and both expressing tRNA partners for the aaRS (Fig. 2b). Target proteins for Tet3Bu incorporation were either a sfGFP150 test protein, or a CRBN^#^ protein, with the superscripts denoting the residue for Tet3Bu incorporation. C-or N-terminal FLAG tags were added to enable identification in western blots, and for testing expression and reaction conditions. For use as a control, an additional version of the second plasmid encodes wild type CRBN (CRBN^WT^).

For the click-chemistry ELF-degrader ligation reagents, we designed six chimeric ―sTCO-JQ1 degraders‖ (Fig. 2c) linking a reactive sTCO group with the JQ1 BRD2/4 ligand ^45, 46^. Linkers ranged from a 2-carbon chain (C2) up to 5 ethylene glycol groups (PEG5) to provide flexibility for how the neo-substrate BRD2/4 was presented to CRBN. In describing these reagents, we use ―JQ1‖ to refer to the bioactive (+)JQ1 stereochemistry and ―(-)JQ1‖ to refer to the inactive enantiomer.

### ELF degrader recruitment of BRD2/4 as a CRBN neosubstrate

In generating CRBN^Tet^ mutants to test for promoting TPD through reaction with sTCO-JQ1 degraders, we first picked five sites around the known IMiD-binding pocket of CRBN (residues 353, 355, 376, 377, 388) thinking that they would best mimic the TPD function of the dBET6 PROTAC molecule, along with one contrasting site (residue 70) that is far both from the IMiD-binding pocket and predicted protein interactions (Fig. 2d).

#### Expression and high-yield in-cell reactivity of containing CRBN^Tet^ forms

To test the ability of this GCE system to make Tet-containing proteins in the HEK293T cells, we co-transfected cells with the Tet3Bu-RS plasmid along with a plasmid encoding either sfGFP150, a CRBN^Tet^ form or CRBN^WT^ (Fig. 2b). To confirm the expression and reactivity of the Tet-containing proteins, we cultured the cells in the presence and absence of Tet3Bu, lysed the cells and reacted some lysate with sTCO-PEG_5K_ (Fig. 3a,b). These results demonstrated the robustness of the expression system, with full-length expression of the sfGFP control and the CRBN^Tet^ forms dependent on the presence of Tet3Bu, and all CRBN^Tet^ forms expressed at levels similar to that of CRBN^WT^. Also, for the Tet-containing proteins, full reactivity was evident, as >95% of each protein was shifted to a higher molecular weight, whereas the control CRBN^WT^ control remained unshifted (Fig. 3b). Based on that success, we carried out the same tests in HEK293T CRBN knockout (K/O) cells. Again, all proteins expressed well with Tet3Bu incorporated, although with a somewhat greater variation in amounts, and they reacted quantitatively *in vitro* with sTCO-PEG_5K_ (Fig. S1). We conclude that in both cell lines this GCE system gives full incorporation of a reactive Tet ncAA into each CRBN mutant.

**Fig 3:**
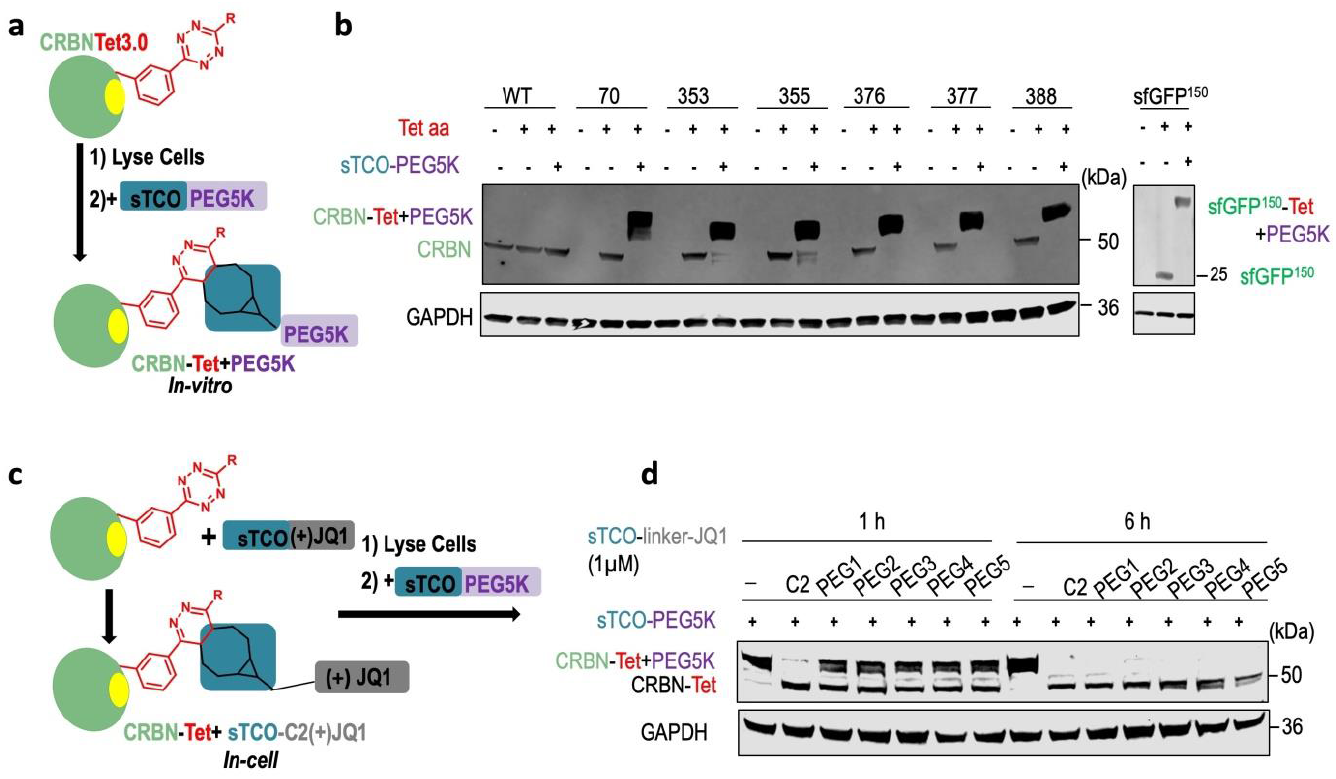
Expression and ligation of CRBN^Tet^ forms in mammalian cells. (a) Design of mobility shift assay for the *in vitro* reactivity of expressed CRBN^Tet^ forms with sTCO-compounds. (b) Annotated FLAG-tag-based immunoblot for CRBN variants (left hand gel) and sfGFP150 control (right hand gel) documenting expression levels in HEK293T cells and reactivity based on mobility shift after *in vitro* reaction with sTCO-PEG_5K_. GAPDH loading level control included. (c) Design of mobility shift assay for the *in vivo* reactivity of CRBN^Tet^ forms with sTCO-JQ1 degraders. (d) Annotated immunoblot, as in Panel b, of mobility shift assay to detect residual reactivity of CRBN^355^ after *in vivo* reaction for either 1 or 6 h with 1 μM of each of the six sTCO-JQ1 degraders.

Next, before moving into mammalian cells, we reacted the sTCO-C2-JQ1 degrader *in vitro* with purified Tet-containing sfGFP150 to test the ability of the degraders to ligate Tet-containing proteins. Importantly, since reaction with a sTCO-JQ1 degrader will block a later reaction with sTCO-PEG_5K_, a simple assay we use throughout this study for the completeness of reaction with a degrader is the lack of a mobility shift upon subsequent reaction with sTCO-PEG_5K_. This mobility shift assay showed that a 15-minute reaction using 3-equivalents of the sTCO-C2-JQ1 degrader led to complete labeling (Fig. S2a), and the integrity of the product was confirmed by mass spectrometry (Fig. S2b).

To achieve quantitative in-cell labeling of Tet-proteins using the sTCO-JQ1 degraders, we used CRBN^355^ as the test protein, and sought conditions which fully blocked the subsequent reaction – after cell lysis – with sTCO-PEG_5K_ (Fig. 3c,d). Trying degrader concentrations of 0.01, 0.1 and 1 μM and incubation times of 1 h and 6 h, we found that only a 6 h incubation at 1 μM of sTCO-JQ1 degrader gave complete labeling of CRBN^355^, and that these conditions worked across all linker lengths (Fig. 3d). At 0.01 and 0.1 μM degrader concentrations, the fraction of CRBN^Tet^ labeled appeared to be roughly 10% and 50%, respectively (Fig. S3).

#### ELF degraders applied to multiple CRBN^Tet^ forms direct the degradation of BRD2/4

Since CRBN^Tet^ expression requires the simultaneous transient transfection of two plasmids, a subset of cells will not express any CRBN^Tet^ protein, and thus will not exhibit any ELF degrader-dependent TPD. To avoid considering these cells when quantifying TPD in the transiently transfected CRBN-knockout (KO) cells, we used a flow cytometer based-immunofluorescence (IF) assay. In this assay we identified CRBN-positive cells using an anti-FLAG antibody, and then used signals from anti-BRD2 and BRD4 antibodies to quantify those proteins in the CRBN-positive cells. The fraction of cells expressing CRBN was generally in the 30 - 60% range (Fig. S4). To establish the validity of the IF assay, we carried out a classic PROTAC assay using dBET6 treatment of cells transiently expressing FLAG-tagged CRBN^WT^ and measured BRD2/4 levels by the IF assay (Fig. 4a). Consistent with what has been seen by others ^50^, the assay showed BRD2 and BRD4 degradation of 90% and 80%, respectively, and control compounds such as JQ1 by itself and its inactive (-)JQ1 isomer led to no BRD2/4 degradation.

**Fig 4:**
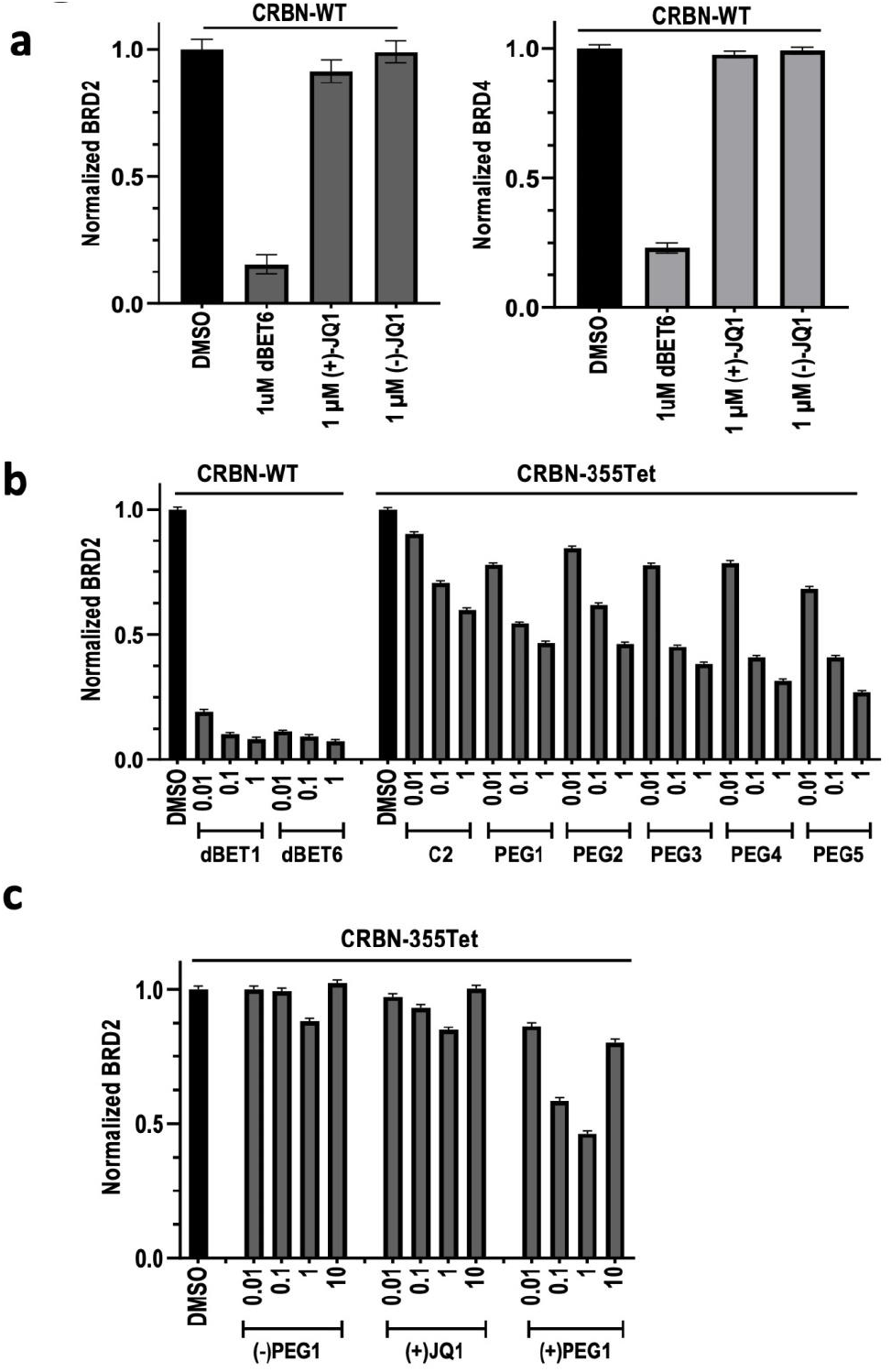
Ligation of CRBN^355^ with ELF degraders promotes degradation of BRD2. All panels show the fraction of remaining target protein (BRD2 or BRD4) as a function of treatment, noted on the x-axis. All IF assays were carried out using HEK293T CRBN K/O cells transfected with 800 ng CRBN^WT^ or CRBN^355^ plasmid and grown at least 36 h. (a) Cells expressing CRBN^WT^ were treated with 1 μM of dBET6, active (+)JQ1, or inactive (-)JQ1 for 6 h before assaying. (b) CRBN^WT^ control results (left hand conditions) are shown with CRBN^355^ results from cells grown with 30 μM Tet3Bu then treated for 6 h with 0.01, 0.1, 1 μM of sTCO-JQ1 degraders. (c) CRBN^355^ negative controls using (-) PEG1 (a degrader with the inactive (-) JQ1 isomer) and JQ1 alone. Also included are CRBN^355^ results using the PEG1 degrader at 0.01, 0.1, 1 and 10 μM.

Next, we evaluated the effectiveness of the full set of six sTCO-JQ1 degraders using one target protein, BRD2, and one CRBN^Tet^ form, CRBN^355^. We chose residue 355 because it is right at the dBET6 binding pocket, and we hypothesized that CRBN^355^ when conjugated to sTCO-JQ1 degraders would best mimic dBET6 bound to CRBN^WT^ and best enable BRD2 degradation. Further controls transfected with CRBN^WT^ and exposed to 0.01, 0.1 and 1.0 μM dBET1 and dBET6 all showed ∼90% degradation of BRD2, and the cells containing CRBN^355^ exposed to ELF degraders over the same concentration range showed a dose-dependence reduction in BRD2 for all degraders, with a maximal value of 75% degradation for the sTCO-PEG5-JQ1 degrader at 1.0 μM (Fig. 4b). Additionally, a clear trend of increased BRD2 degradation was observed with increased ELF degrader linker length. The dose dependent increase makes sense given that the ligation of the CRBN^Tet^ increases with degrader concentration and only reaches completeness at 1.0 μM degrader (Fig. 3d and S3). Evidence that BRD2 degradation is induced by the proposed ELF degrader mechanism, is provided by negative controls showing that BRD2 degradation is only minimally stimulated by JQ1 itself or a degrader containing the inactive (-)JQ1 isomer (Fig. 4c). Finally, tests of degraders at a higher 10 μM concentration resulted in much less degradation (Fig. 4c and S5), a behavior also generally observed in PROTAC studies and referred to as the ―hook effect‖ ^51, 52^. The explanation for this decrease is that excess degrader (i.e. degrader not ligated to CRBN^Tet^) can bind to the neosubstrate and block its binding to the degraders that are ligated to CRBN^Tet^.

Given the above success with CRBN^355^ and BRD2 degradation, we expanded our study of ELF degraders to monitor the degradation of both BRD2 and BRD4, and evaluate all of the initially selected six Tet sites on CRBN, five around the known IMiD-binding pocket of CRBN and one (residue 70) rather distant (Fig. 2d, 5a). Also, given the clear trends with ELF degrader length and concentration, in these further studies, we only tested the shortest and longest PEG-linked degraders (PEG1 and PEG5 respectively), and tested them only at the concentrations of 0.1 and 1.0 μM. At all six sites the ELF degraders promoted the degradation of both BRD2 and BRD4 and the dependencies on the degrader length and concentration matched those seen in the pilot studies, with the longer linker and the 1.0 μM concentration being more effective in every case (Fig. 5b,c). Even though all CRBN sites promoted degradation, the sites near the IMiD-binding pocket were much more effective, leading to a high 50-75% level of degradation of both BRD2 and BRD4, whereas site 70, far from the IMiD-binding pocket, led to just a 40% degradation of BRD2 and an even lower 20% degradation of BRD4. While this indicates that different surfaces of CRBN are differentially effective in promoting degradation, and that ELF degrader ligation of CRBN^70^ promotes degradation at all indicates that there is substantial structural plasticity in how CRBN can recruit substrates.

**Fig 5:**
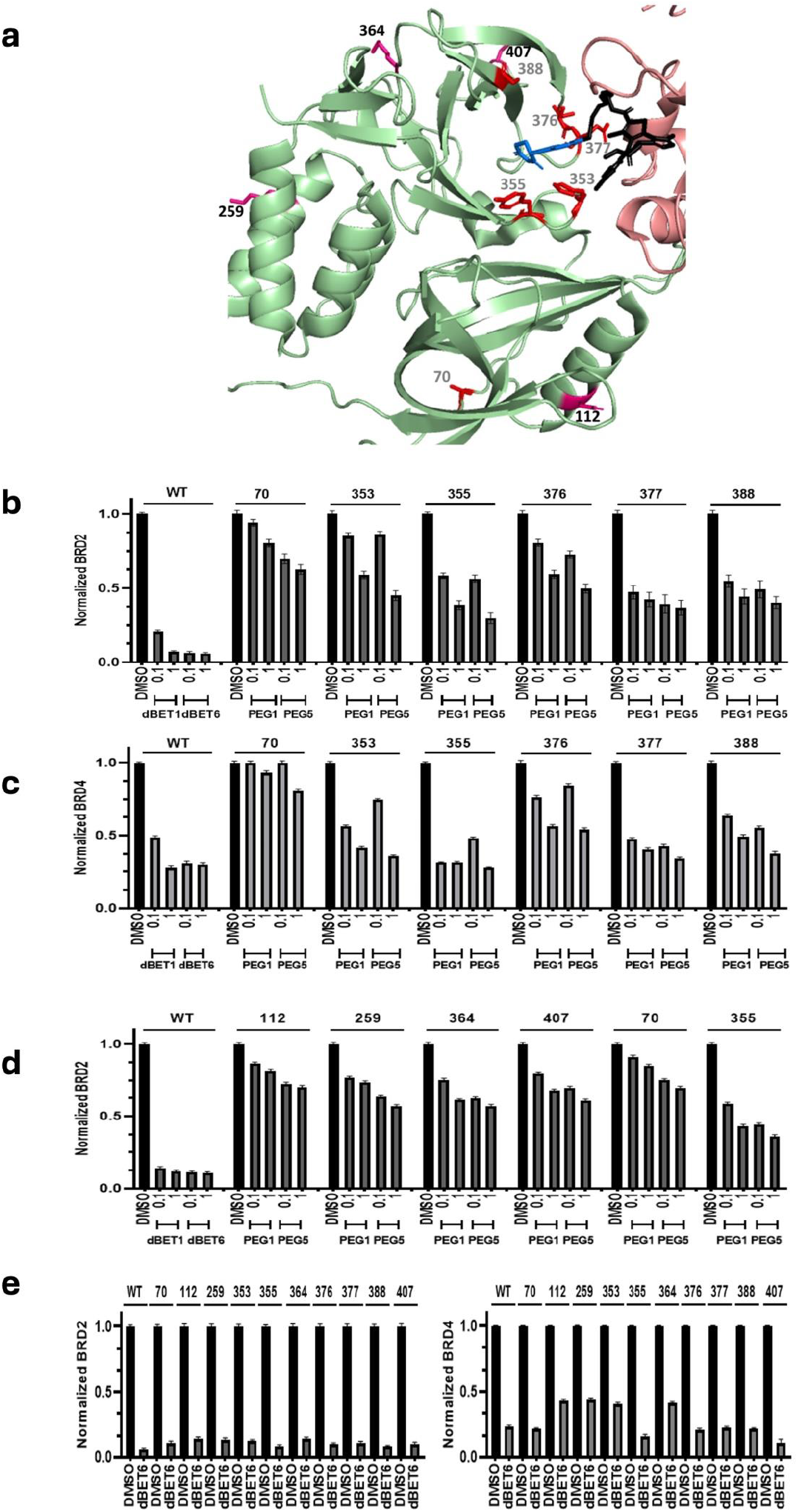
ELF degrader effectiveness varies as a function of the CRBN surface position. IF assay result plots are as in Fig. 4. (a) Ribbon diagram of CRBN: BRD4 complex as in Figure 2d, but showing side chains (red sticks with residue numbers) for the six originally targeted Tet sites (grey residue numbers) and the four additional ones (black residue numbers). (b) BRD2 degradation results for original CRBN^Tet^ target sites using 0.1 and 1 μM concentrations of the PEG1 and PEG5 degraders. Positive CRBN^WT^ controls with dBET1 and dBET6 and a negative DMSO solvent control are included. (c) Same as Panel b, but for BRD4 degradation. (d) Same as Panel b, but for four additional CRBN surface sites (112, 259, 364, 407) shown in Panel a. (e) IF results for BRD2 (left-hand plot) and BRD4 (right-hand plot) degradation promoted by 1 μM dBET6 treatment of each of the 10 CRBN^Tet^ forms.

#### Additional probing of the structural plasticity of CRBN promoted TPD

Given that the promotion of BRD2/4 degradation by ELF degraders was not limited to ligating CRBN residues at or near the IMiD binding pocket, we further probed the structural plasticity of CRBN function by picking four additional sites (residues 112, 259, 364 and 407) that sample the addition exposed surface areas of CRBN (Fig. 5a). Except for CRBN^112^, the new CRBN forms showed Tet-ncAA dependent CRBN^Tet^ expression and robust in cell labeling with 1 μM of the sTCO-PEG5-JQ1 degrader (Fig. S6); for unknown reasons, CRBN^112^ showed leaky expression in the absence of Tet ncAA, but a substantial fraction of the protein underwent in cell labeling with 1 μM ELF degrader, meaning it was suitable for study. Remarkably, all of these additional CRBN^Tet^ forms showed a substantial ELF degrader effects, inducing 30-40% degradation of BRD2 (Fig. 5d). Interestingly, CRBN^112^ and CRBN^70^ which are on the same face of CRBN, both showed a slightly lower ∼25% degradation of BRD2, while CRBN^Tet^ sites 259, 364 and 407, which are spread out on a different face of CRBN (Fig. 5a), led to a slightly higher degradation of ∼40%.

#### Tet-encoding is generally compatible with PROTAC function

A final question we asked is the extent to which the CRBN^Tet^ protein forms can still participate in PROTAC stimulated TPD. To test this, we took the full panel of CRBN^Tet^ forms and assessed the ability of dBET6 to stimulate the degradation of BRD2 and BRD4.

The levels of BRD2 and BRD4 degradation achieved by CRBN^WT^ were ∼95% and ∼75%, respectively. For both BRD2 and BRD4 none of the CRBN^Tet^ forms showed substantial impairment, with all achieving degradation levels at least 80% of those stimulated by the wild-type protein (Fig. 5e). Also, the extents of impairment shown by the various forms were reasonably consistent for the two protein targets, implying that the mechanisms of impairment are the same for the two substrates. The CRBN^Tet^ forms with Tet at sites 112, 259, 353 and 364 showing the most impairment for both protein targets, with the degradation being ∼85% and ∼60% for BRD2 and ∼BRD4, respectively.

## DISCUSSION

The discovery that TPD could be accomplished by repurposing the cell’s UPS has revolutionized therapeutic strategies by providing access to the massive intracellular undruggable proteome ^53^. However, even though many PROTAC and ―molecular glue‖ therapeutics have been developed, the huge potential for application to degrade specific, disease-causing target proteins, has been limited due to the small number of E3 ligases ^33, 34, 35^ for which high-affinity ligands are known ^36, 37^. Indeed, even while computational methods can identify potential sites for ligand development on novel E3 ligases, no methods exist to evaluate whether a PROTAC-like molecule binding at the putative ligand site would also have a geometry that could drive target protein ubiquitination and subsequent degradation. It is to address this gap in knowledge that we sought here to develop an E3 ligase ligand free degrader (ELF-degrader) that can be site-specifically positioned on any E3 ligase in live cells to map the surface of E3 ligase and identify optimal pockets for TPD.

Genetic code expansion (GCE) technology offers a powerful tool for a minimally-perturbing site-specific incorporation of new chemical functionality into proteins in live cells, and this enables protein interactions to be studied in their native cellular environment. While GCE has been use to covalently identify interacting proteins with photo-reactive ncAAs to identify protein interactions and evaluate protein dynamics in cellular processes ^54, 55, 56^, encoding bioorthogonal chemistry was selected to provide high labeling efficiencies in short times ^44, 57, 58, 59^. While many GCE encoded chemistries have enabled antibodies functionalization to form antibody drug conjugates, few have the labeling efficiency or speed to quantitatively label proteins for functional alteration in live cells. Here in this work, we choose to encode tetrazine amino acids on the E3 ligase surface because this approach has the fastest in cell labeling rates with sTCO functionalized ligands^60,61^. We selected the CRBN E3 ligase to identify the limits of the ELF-degrader system, since Tet-ncAAs encoded near the IMiD-binding pocket could be labeled with sTCO functionalized with JQ1 moiety and mimic dBET6 (a potent and selective degrader that comprises JQ1 conjugated to a CRBN specific binder pomalidomide) brings the targeted BRD domain in proximity with the ubiquitin ligase complex and lead to efficient ubiquitination and proteasomal degradation. By altering the surface locations of the Tet-ncAA and the nature of the linker composition of sTCO-JQ1 degrader molecules (Fig. 2), we can monitor cellular levels of BRD2 and BRD4 and optimize ELF-degrader conditions.

First, we picked five sites (Fig. 2d) around the IMiD-binding pocket to encode Tet3Bu and along with one control site 70 far from predicted protein interactions and the IMiD-binding pocket. For all sites we demonstrated complete Tet3Bu encoding and their ability to label quantitively in HEK293T CRBN K/O cells confirming the robustness of this GCE approach (Fig 3). Next we confirmed that all the sTCO-JQ1 degraders with varying linker composition could enter cells and quantitatively label the CRBN^Tet^ (Fig 3d). This ability allowed us to proceed with evaluating a variety of CRBN surface attachment sites and degrader linker lengths in terms of how well the resulting CRBN-JQ1 conjugates degraded protein targets directly in live cells.

To monitor TPD, we developed an IF staining assay which specifically monitors the cellular level of the BRD2/4 target proteins in E3 ligase producing cells and validated the assay by demonstrating 75-95% loss of BRD2/4 levels in the presence of dBET1 and dBET6, while control compounds such as JQ1 by itself and its inactive (-)JQ1 isomer led to no BRD2/4 degradation (Fig 4 a,b). To evaluate the effectiveness of the six sTCO-JQ1 degraders at low concentrations (0.01 to 1.0 μM), we monitored the loss of BRD2 target when cells containing CRBN^355^, a variant placing the Tet-ncAA right at the edge of the dBET6 binding pocket (Fig 4b). This clearly demonstrated that ELF degraders would work, as at the optimal degrader concentration of 1 μM and with the longest of the linkers (PEG5), the cellular BRD2 was reduced by 75%. Minimal degradation of BRD2 occurred when cells are exposed to JQ1 itself or a degrader containing the inactive (-) JQ1 isomer (Fig. 4c), supporting our proposed ELF degrader mechanism. As expected substantially less target degradation was detected at a higher 10 μM degrader concentration, a behavior referred to as the ―hook effect‖ in PROTAC studies^51, 52^. A primary reason for selecting a GCE labeling system with fast intracellular labeling rates was to enable avoiding the ―hook effect‖ by using low label concentrations such that the ligation could proceed in a reasonable timeframe, even while having a minimal excess of unreacted label.

This validation of ELF degrader function encouraged us to evaluate the robustness of this approach for removing BRD2 and BRD4 using the four other CRBN^Tet^ sites around the IMiD-binding pocket and the CRBN^70^ site far from IMiD-binding pocket (Fig 2d, 5a). Impressively all sites near the IMiD-binding pocket, after reaction with an ELF degrader, showed 50-75% degradation of both targets when the longer PEG5 linker was used and slightly less degradation with the shorter PEG1 linker (Fig 5b,c). And interestingly, these five sites led to nearly equivalent (within ∼5% of each other) degradation levels of BRD2 and BRD4. In contrast, ligation at the distant CRBN^70^ site promoted degradation to a lesser extent, degrading ∼40% of BRD4 and ∼20% of BRD2. This is an important result, because it indicates that there is substantial structural plasticity in how CRBN can recruit targets, while also making clear that surface site location can lead to functional preferences that differentially impact different target proteins.

To further evaluate the structural plasticity of CRBN with ELF degraders, we tested four additional sites that sample additional exposed surfaces of CRBN. We discovered that the two sites (70 and 112) on the same face but far from the IMiD-binding pocket had similarly low BRD2 degradation ability (10-25%) whereas the three CRBN^Tet^ sites 259, 364 and 407, on the opposite face showed improved (25-40%) degradation ability (Fig. 5b, d). This ability to ―walk the ELF degrader‖ position around the surface of E3 ligases and with site-specific control probe functional constraints and potential ligand binding pockets reveals how the ELF degrader approach is a powerful tool that makes it possible to efficiently explore the therapeutic potential of E3 ligases for which ligands have not been identified.

Our final experiment was motivated by the realization that even though Tet-ncAA encoding on an E3 ligase is a small structural change, it could still compromise protein conformation or stability and negatively impact CRL4-complex formation or dBET6 binding and thus function. Since all CRBN^Tet^ forms enabled BRD2/4 degradation at levels on par with those stimulated by CRBN^WT^ (Fig 5e), we conclude that Tet-modified E3 ligase forms will, in general, be compatible with PROTAC function. Interestingly, of the four most impaired sites, only one (residue 353) was at the PROTAC binding site where it is easy to understand how the mutation could impact PROTAC binding. The other three (residues 112, 259 and 364) were at surface positions rather distant from the PROTAC binding site (Fig. 5a), implying that the mutations at these sites are somehow impacting the assembly or functionality of the CRL4-complex.

A modeling of open and closed CRL4-complexes with ELF degraders attached at different CRBN^Tet^ surface positions (Fig. 6) shows how the ELF degrader results are consistent with a simple distance model for how the site of attachment alters the extent of target protein degradation. The CRBN^355^ position modified with the longest (the PEG5) ELF degrader and bound to BRD4 shows a very short distance for ubiquitin transfer for both the open and closed complex forms. Alternatively, CRBN^70^ when bound to BRD4 has a much longer distance for ubiquitin transfer in both open and closed forms. And the degradation data correlates well with distance, as attachment sites around the IMiD-binding pocket produce the highest degradation levels, sites with the longest distances (70 and 112) show the lowest degradation levels, and sites with a middle distance (359, 364, and 407) provide a medium degradation level. This modeling also shows that the conformational shift from open to closed CRBN forms does impact the geometry for ubiquitination (Fig. 6a versus b). Notably, since ELF degrader ligation would not be expected to trigger the conformational change, the lack of such a change could be a factor in why none of the CRBN^Tet^ forms ligated with ELF degraders achieved the same level of BRD2 degradation of as is promoted by dBET1 or dBET6.

**Fig 6:**
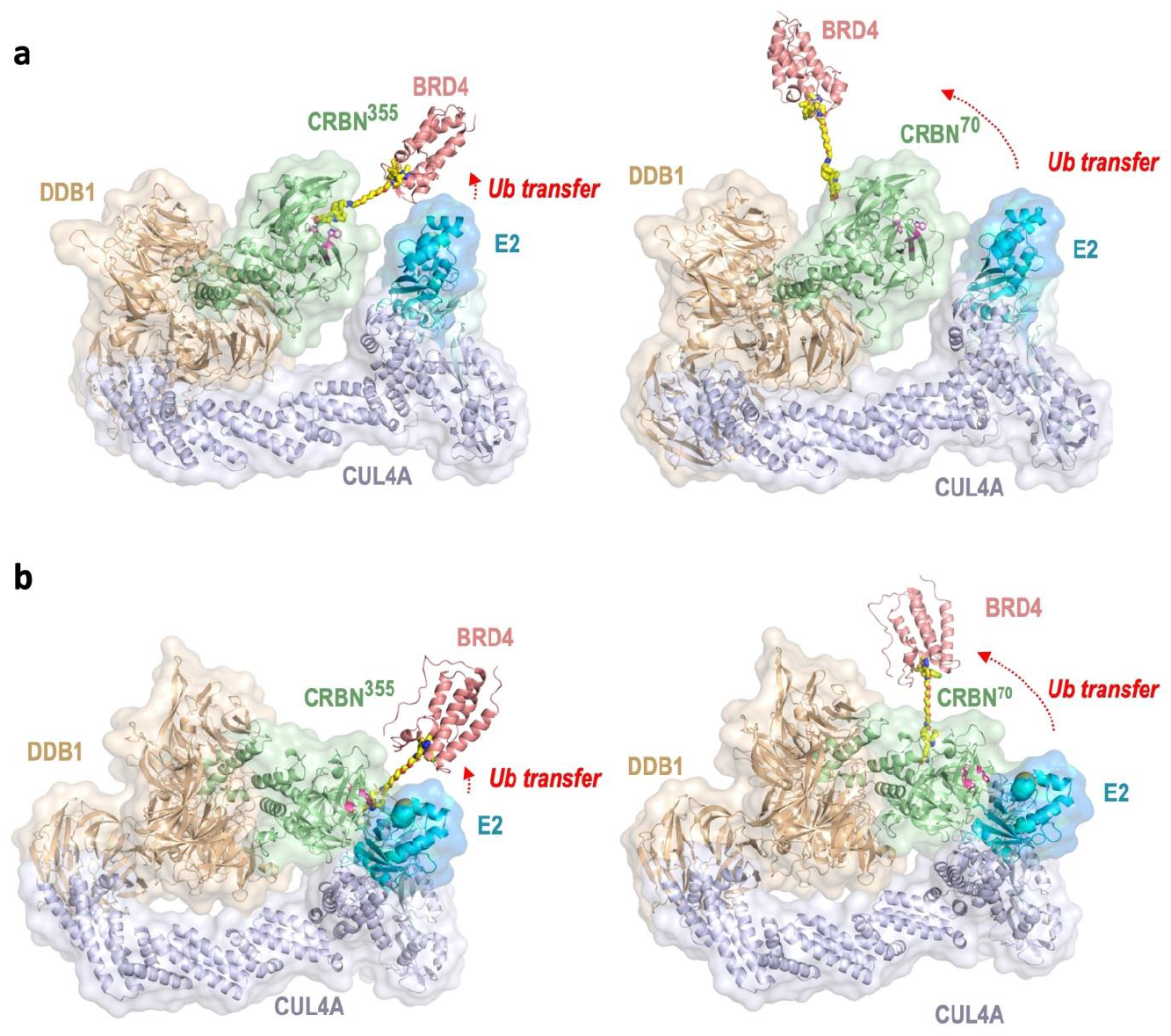
Modeling supports a distance-based explanation for ELF degrader effectiveness. (a) The open conformation of a CRL4 complex including CRBN and BRD4 is shown, with the BRD4 molecule moved to be placed at the end of a modeled PEG5 linker attached at residue 355 (left hand panel) or 70 (right-hand panel). Transparent surfaces and ribbon diagrams for each protein (colored as in Fig. 1b) are labeled and the distance over which ubiquitin must be transferred is highlighted (red text and arrow). (b) Same as Panel a but based on the closed conformation of a CRL4 complex including CRBN and BRD4. The core model consisting of E2 (UBE2G1, uniprot no. P62253), RBX1 (uniprot no. P62877), CUL4A (uniprot no. Q13619) and the BPB domain of DDB1 (residues 393-709, uniprot no. Q16531) were generated using AlphaFold3 ^62^. The open and closed conformations were generated by overlaying the DDB1 BPB domains from the twisted (PDB 2HYE,^63^) and hinged (PDB 5HXB,^64^) CRBN/DDBP1 structures onto the DDB1 BPB domain of the AlphaFold3 generated core structure, respectively. The JQ1 moiety of the JQ1-PEG5-sTCO ELF degrader (generated using Phenix Elbow,^65^) was docked into BRD4 by overlaying it with the same moiety of the BRD4/JQ1 co-complex structure (PDB 3MXF,^66^).

In summary, we demonstrate that the site specific encoding of Tet-ncAAs can be used to rapidly functionalize E3-ligases in live cells with protein-targeted degrader ligands and that this robust system can be used to evaluate the structural organization of the UPS. The design of the ELF-degrader approach enables it to be applied to any E3 ligases for exploring not only their potential for neo-substrate TPD, but also to specifically explore those sites on their surfaces that are computationally identified as attractive sites for ligand development. Furthermore, since we showed this GCE bioconjugation approach of encoding Tet-ncAA on CRBN is compatible with traditional PROTAC function, this work also opens the door to using on-protein Tet ligation to attach a fluorescent probe to functional E3 ligases and track changes in complexation and sub-cellular localization under native states and in response to PROTAC treatment.

## METHODS

### Plasmids

All the plasmids used for the experiments have been summarized in Table 1. The synthetase plasmid was pAcBac1-R2-84-RS, pAcBac1-sfGFP-150TAG (Plasmid Control for Tet expression having C-term FLAG tag), pAcBac1-sfGFP-WT (transfection control plasmid), Gene of interest (GOI) plasmids were pAcBac1-CRBNWT (N-term FLAG tag), pAcBac1-CRBNTAGs (Table SI-1).

### Transfection of HEK cells

HEK293T and HEK293T CRBN K/O cells were grown in DMEM /10% FBS (Fetal Bovine Serum). The cells were plated in a 24-well plate at 1:3 ratios from the 100 % confluent plate so that they reach 70% to 80% confluency at the time of transfection. Cells were transiently transfected using JetPrime (***Polyplus***-***transfection***®) using the manufacturer’s protocol with minor modification. Briefly, 600-800 ng of plasmid DNA was taken in 1:8 [RS: GOI], 50 μl of JetPrime Buffer, and 1.2 μl of JetPrime reagent. They were combined and incubated for 20 min at RT before adding to cells. Tet3Bu amino acid at 30 μM was added to the cells immediately and incubated for 24-48 h, depending on the experiment, before analysis.

### *In-Vivo* Tetrazine labeling and mobility shift assay

To examine the specific *in-cell* labeling of CRBN^Tet^ proteins in HEK293T cells, transfected cells were washed two times with DMEM media and incubated for 1 or 6 h with 1 and 10 μM sTCO-Jq1 degraders. Unreacted degraders were quenched by at least three molar excess of Tet2-Me ^49^. Cells were washed with PBS and lysed using RIPA lysis buffer system (Santa Cruz Biotechnology, sc-24948). To the cell lysate, 20μM of sTCO-PEG_5K_ was added for 15 min and then quenched with Tet-v2.0Me for 5 min. The samples were then prepared using β-mercaptoethanol Laemmli buffer and after thermal denaturation, the lysates were separated by 12% SDS-PAGE followed by transfer to nitrocellulose membranes.

The membranes were blocked with Licor Blocking buffer for 60 min and then washed three times with TBS-T (tris-buffered saline containing 0.1% Tween-20), incubated with a primary (Mouse AntiFLAGM2) antibody for 60 min, washed three times with TBS-T and then incubated for 60 min with a secondary (Anti Mouse 680IR) antibody followed by washing again three times with TBS-T before imaging. Immunoreactive bands were detected Licor Instrument.

### The immunofluorescence (IF) assay for assessing CRBN-mediated degradation of target protein (BRD2/4)

HEK293T or CRBN-KO cells were transfected with 800 ng of CRBN^WT^ or CRBN^Tet^-expressing plasmid DNA, and grown for at 36 – 48 h, in the case of CRBN^Tet^-expressing cells, 30 μM Tet3Bu was present. Cells then underwent treatment for 6 h with a TPD agent (either dBET6, dBET1 or an ELF degrader) or a control compound. After treatment, cells were prepared using BD cytofix/cytoperm fixation/permeabilization kit (BD Biosciences) according to the manufacturer’s instructions. Cells were first stained with FLAG primary antibody (Sigma F1804, 1to200 dilution), and then stained for BRD2 (Abcam ab197865, 1to50 dilution), anti-mouse IgG (Thermofisher A-32742, 1to200), and BRD4 (Abcam ab197608, 1to50). All staining procedures were performed for 30 min at 4 °C. Samples were analyzed on an LSR Fortessa X-20 (BD Biosciences), and data were processed using FlowJo (v.10.10),0) analysis software (BD Biosciences). FLAG-positive singlets were manually gated, and the population median fluorescent intensity (MFI) and median absolute deviation (MAD) were extracted from each condition for visualization. In every acquisition at least 1000 cells were collected in the final FLAG + population, it’s the variation within the cell population.

## Acknowledgments

We are grateful for the assistance from Jeff Moore for mass spectrometry data acquisition and processing.

## Funding

This work was supported in part by the GCE4All Biomedical Technology Development and Dissemination Center supported by National Institute of General Medical Science grant RM1-GM144227 and 1S10RR025628-01 to the Oregon State University Mass Spectrometry Facility.

## COMPETING INTERESTS

The authors declare the following competing financial interest(s): Y.Z., Z.N., V.M., J.Y., S.W., A.V., J.M.R. are current employees of AbbVie. P.R. was an employee of AbbVie at the time of the study. The design, study conduct, and financial support for this research were provided in part by AbbVie. AbbVie participated in the interpretation of data, review, and approval of the publication.

