## Supplemental for "An in-cell approach to evaluate E3 ligases for use in targeted protein degradation": Supplementary File.docx

**Table of Contents**

1. **Supplemental Methods	pages S1-S8**
2. **Supplemental Figures	pages S9-S13**
3. **Supplemental Tables	pages S14**
4. **^1^H and ^13^C NMR Spectra	pages S15-S30**

**sfGFPWT and sfGFP150-Tetv3.0 Butyl protein expression and purification:** A cell stock of BL21(DE3) cells co-transformed with the Mb-pAJE-Tet3.0 (GCE plasmid), and pET28-sfGFP-wt (GOI) that expresses sfGFP wt control protein with C-terminal His6 tag, under a T7 transcriptional promoter, lac operon and pET28-sfGFP150TAG (GOI). The transformed colony was used to inoculate a 5 mL culture of non-induction media (NIM) containing kanamycin (50 μg/mL) and spectinomycin (100 μg/mL), which was then grown 16 hours at 37 °C shaking at 250 rpm. A 50 mL auto-induction media (AIM) culture containing kanamycin (50 μg/mL) and spectinomycin (100 μg/mL) was inoculated with 0.5 mL of the grown NIM. The AIM was supplemented with 0.5 mM Tet- v3.0 from a 100 mM DMF stock solution. The AIM culture was allowed to grow shaking at 250 rpm for 24 hours at 37 °C. Additionally, cultures without Tet-v3.0 and containing only the WT plasmid were grown simultaneously. After 24 hours OD_600_ was measured, and for control cultures fluorescence of sfGFP and sfGFP-150TAG (ex/em: 485/510nm) was measured. As expected, sfGFP-wt was visibly green while sfGFP-150TAG was orange due to quenching by the incorporated Tet ncAA. The OD_600_ values for sfGFPWT was 2.1 and sfGFP150 was 2.5. All cells were harvested by centrifugation at 5000 rcf for 5 min. Supernatant was decanted and cell pellets were stored at -80 °C.

To purify the sfGFP forms, cells were resuspended in lysis buffer (50 mM Na_2_PO_4_ pH 7.0, 500 mM NaCl, 5 mM imidazole, pH 7.0), lysed using a Microfluidics M-110P microfluidizer (18,000 psi), and the collected lysate clarified by centrifugation (21000 rcf, 1 h). To the decanted supernatant, we added 800 μL bed volume TALON resin, and incubated the mix for 1-2 h gently rocking at 4° C. After pouring a column with the resin-lysate slurry, discarding the flow through, the resin was washed with 3 x 10 mL lysis buffer. Protein was eluted with 2.5 ml of elution buffer (50 mM Na_2_PO_4,_ 500 mM NaCl, 250 mM imidazole, pH 7.0), and the eluate was desalted using a PD-10 column (GE Healthcare) with 3.5ml of the desalting buffer (50 mM Na_2_PO_4,_ 100 mM NaCl, pH = 7.0). The purity of protein was verified by gel electrophoresis and its concentration measured by absorbance at 280 nm (Ɛ = 24080 M^−1^ cm^−1^ for GFPWT and Ɛ = 35734 M^−1^ cm^−1^ for GFP150TAG). Protein samples having 3mg/ml and 1.7 mg/ml concentrations were aliquoted and stored at −80 ℃.

**Testing Tet-v3.0 Protein Reactivity Using Mobility Shift Assay**

Purified protein forms were allowed to stand for overnight to assaying reactivity to ensure the Tet3Bu was fully oxidized (and thus become fully reactive). To the purified protein (8 µg), a 10 molar excess of sTCO-PEG_5K_ was added and incubated 15 min at room temperature followed by quenching with a 12 molar excess of Tet2-Me 5 min, RT. The samples were then prepared using β-mercaptoethanol Laemmli buffer and after thermal denaturation and ran onto the 12 % SDS-PAGE.

**Mass spectral analysis of sfGFP-Tet-3.0**

Purified desalted sfGFP-TAG150-Tet3Bu (50 µM) was reacted with 2 or 3 equivalents of sTCO-C2-JQ1 for 15 min at room temperature and then further desalted using NAP-5 Columns with LC-MS grade water to a final concentration of 30-50 µM using a 2 ml, 10 kDa MWCO centrifugal filter (Millipore). The samples were further analyzed using an FT LTQ mass spectrometer at the Oregon State University mass spectrometry facility.

**sTCO-JQ1-ELF Degraders Synthesis**

*Reagents:*

Unless otherwise specified all the reagents and solvents were obtained from Sigma-Aldrich (Milwaukee, WI). (+)JQ1 carboxylic acid was purchased from Combi-Blocks (San Diego, CA). (-)JQ1 carboxylic acid was purchased from Enamine US. All PEG linkers reagents were purchased from Broadpharm (San Diego, CA). Anhydrous solvents were purchased from Sigma-Aldrich in sealed bottles (Sure/Seal™).

*Analytical Methods*

Crude reaction monitoring and final product purity checks were performed on a waters Acquity UPLC system quipped with an in-line photodiode array detector (PDA) and SQD mass spectrometer, running MassLynx 4.1 and Openlynx 4.1 software (Waters Corporation, Milford, MA, USA). The SQD mass spectrometer was operated under positive ESI ionization conditions. The column used was a Waters BEH C8, 1.7μm (2.1mm × 30mm) at a temperature of 55°C. The following ammonium acetate analytical method was used: a gradient of 10- 100% acetonitrile (A) and 10 mM ammonium acetate in water (B) was used, at a flow rate of 1.0 mL/min (0-0.1 min 10% A, 0.1-1.1 min 10-100% A, 1.1-1.3 min 100% A, 1.3-1.4 min 100-10% A). NMR spectra were acquired at 27 ^o^C on various spectrometers (Varian or Bruker) equipped with sample changers. Chemical shifts (ppm) are given relative to solvent: references for DMSO-*d*_6_ were 2.50 ppm (^1^H NMR) and 39.52 ppm (^13^C NMR). NMR data are represented as follows: chemical shift, multiplicity (s = singlet, br s = broad singlet, d = doublet, dd = doublet of doublet, t = triplet, q = quartet, m = multiplet, etc.), coupling constant, and integration. Coupling constants (J) are given in Hertz (Hz). High-resolution mass spectrometry characterization was acquired using a Thermo Scientific™ Orbitrap ID-X™ TribridTM mass spectrometer. Calculated masses listed herein represent monoisotopic masses, [M+H]+ .

Isolation of products from reaction mixtures was accomplished by preparative-scale reverse-phase HPLC on a Gilson HPLC system equipped with liquid handler (Gilson-215) and UV/Vis detector. Separations were performed on a Phenomenex Luna C8 15 µm, 100 Å (225 x 30mm) or a Phenomenex Gemini NX-C18 15 µm, 100 Å (200 x 25 mm) column. An ammonium hydroxide or ammonium acetate purification method was used: a gradient of acetonitrile (A) and 0.1% ammonium hydroxide in water (B) or 10 µM ammonium acetate in water (B) was used at a flow rate of 50 mL/min (45-75% A/B gradient over 14 min). Samples were injected in 2 mL DMSO. An Agilent 1100 Series Purification system was used, consisting of the following modules: Agilent 1100 Series LC/MSD SL mass spectrometer with API-electrospray source; two Agilent 1100 Series preparative pumps; Agilent 1100 Series isocratic pump; Agilent 1100 Series diode array detector with preparative (0.3 mm) flow cell; Agilent active-splitter, IFC-PAL fraction collector/ autosampler. The make-up pump for the mass spectrometer used 3:1 methanol:water with 0.1% formic acid at a flow rate of 1 mL/min. Fraction collection was automatically triggered when the extracted ion chromatogram (EIC) for the target mass exceeded the threshold specified in the method. The system was controlled using Agilent Chemstation (Rev B.10.03), Agilent A2Prep, and Leap FractPal software, with custom Chemstation macros for data export.

**Chemical Synthesis**

sTCO-acid and final compounds reported here are sensitive to heat and acidic conditions. Samples were handled at room temperature and kept as powder at -20^o^C.

**sTCO-acid (1R,8S,9r,E)-bicyclo[6.1.0]non-4-ene-9-carboxylic acid** was synthesized as reported before in the literature.^1^

**(+/-)JQ1-linker-NH_2_** compounds were synthesized as previously reported in the literature^2^ by amidation of (+/-)JQ1-acid with mono Boc-protected bis-amine linkers in DMF using HATU as the coupling reagent and DIEA as the base. Compounds were purified by HPLC method A and then deprotected using TFA and used directly in the next step.

Compounds **1-8** were synthesized in a library format, at a 0.120 mmol scale (unless otherwise specified), in one step, as shown in the general scheme below:

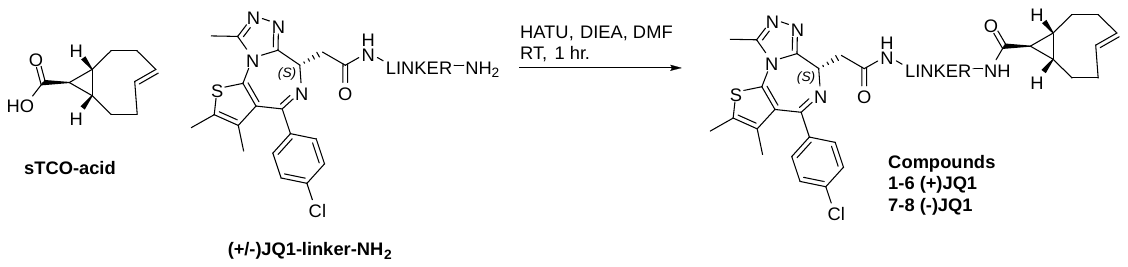

**Scheme SI-1** Synthetic scheme for preparation of **Compounds 1-8**.

**General Procedure A:** sTCO acid (20 mg/reaction, 0.12 mmol, limiting reagent) was activated with 1.1 equiv. of HATU (50 mg, 0.13 mmol) and DIEA (5 equiv. 105 µL, 0.6 mmol) in 1 mL DMF. The reaction was shaken for 5 min at rt, then the resulting solution was added to a 4 mL vial containing amine monomer (1.2 equiv) and the reaction was stirred at room temperature for one hour. Upon completion, reactions were concentrated, dissolved in 2 mL DMSO, filtered, and purified by reverse phase HPLC (ammonium hydroxide or ammonium acetate methods). Fractions containing desired compounds were lyophilized. Compounds were analyzed by LCMS and NMR to assess the % trans:cis.

**Compound 1. (1R,8S,9r,E)-N-(2-(2-((S)-4-(4-chlorophenyl)-2,3,9-trimethyl-6H-thieno[3,2-f][1,2,4]triazolo[4,3-a][1,4]diazepin-6-yl)acetamido)ethyl)bicyclo[6.1.0]non-4-ene-9-carboxamide**

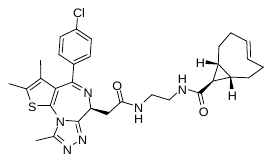

General procedure A was followed using (S)-N-(2-aminoethyl)-2-(4-(4-chlorophenyl)-2,3,9-trimethyl-6H-thieno[3,2-f][1,2,4]triazolo[4,3-a][1,4]diazepin-6-yl)acetamide (64.0 mg, 0.14 mmol) as the amine coupling partner. The crude residue was purified with a Waters HPLC system (mass triggered collection) using a Phenomenex Luna C8 225x30 mm column under ammonium acetate conditions (45-75% CH_3_CN/H_2_O with 10 mM NH_4_OAc, 14 minute run, 50 mL/min), to afford the title compound **1** (7.2 mg, 10 % yield) as a yellow oil. **^1^H NMR** (600 MHz, DMSO-*d*_6_): δ 8.23 (s, 1H), 7.89 (s, 1H), 7.48 (d, *J* = 8.8 Hz, 2H), 7.42 (d, *J* = 8.6 Hz, 2H), 5.80 (ddd, *J* = 16.1, 9.3, 6.2 Hz, 1H), 5.12 (ddd, *J* = 16.8, 10.5, 3.8 Hz, 1H), 4.50 (t, *J* = 5.5 Hz, 1H), 3.23 (dd, *J* = 7.1, 3.7, 2H), 3.17 – 3.11 (m, 4H), 2.59 (s, 3H), 2.41 (s, 3H), 2.30 – 2.11 (m, 4H), 1.94 – 1.82 (m, 2H), 1.62 (s, 3H), 1.06 – 0.98 (m, 1H), 0.95 – 0.82 (m, 3H), 0.64 – 0.54 (m, 1H). **^13^C NMR** (151 MHz, DMSO-*d*_6_) δ 172.6, 169.8, 163.1, 155.1, 149.9, 137.9, 136.8, 135.2, 132.3, 131.4, 130.7, 130.2, 129.9, 129.6, 128.5, 53.8, 38.7, 38.5, 37.7, 37.5, 33.2, 31.4, 28.2, 27.0, 24.2, 24.2, 23.1, 23.0, 14.1, 12.7, 11.3. **HRMS**: (ESI, m/z): calcd. [M+H]^+^ for C_31_H_36_ClN_6_O_2_S^+^: 591.23035, found: 591.23132. trans:cis >20:1 by ^1^H NMR.

**Compound 2. (1R,8S,9r,E)-N-(2-(2-(2-((S)-4-(4-chlorophenyl)-2,3,9-trimethyl-6H-thieno[3,2-f][1,2,4]triazolo[4,3-a][1,4]diazepin-6-yl)acetamido)ethoxy)ethyl)bicyclo[6.1.0]non-4-ene-9-carboxamide**

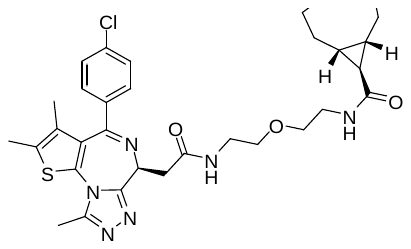

General procedure A was followed using (S)-N-(2-(2-aminoethoxy)ethyl)-2-(4-(4-chlorophenyl)-2,3,9-trimethyl-6H-thieno[3,2-f][1,2,4]triazolo[4,3-a][1,4]diazepin-6-yl)acetamide (70 mg, 0.14 mmol) as the amine coupling partner. The crude residue was purified with a Waters HPLC system (mass triggered collection) using a Phenomenex Gemini NX-C18 225x30 mm column under ammonium hydroxide conditions (45-75% CH_3_CN/H_2_O with 0.1% NH_4_OH, 14 minute run, 50 mL/min), to afford the title compound **2** (33 mg, 43 % yield) as a yellow oil. **^1^H NMR** (500 MHz, DMSO-*d*_6_) δ 8.26 (t, *J* = 5.6 Hz, 1H), 7.92 (t, *J* = 5.5 Hz, 1H), 7.48 (d, *J* = 8.9 Hz, 2H), 7.42 (d, *J* = 8.6 Hz, 2H), 5.78 (ddd, *J* = 16.8, 9.2, 6.2 Hz, 1H), 5.08 (ddd, *J* = 16.7, 10.5, 3.8 Hz, 1H), 4.50 (dd, *J* = 8.3, 5.9 Hz, 1H), 3.48 – 3.41 (m, 4H), 3.30 – 3.17 (m, 6H),f 2.59 (s, 3H), 2.41 (s, 3H), 2.27 – 2.07 (m, 4H), 1.93 – 1.79 (m, 2H), 1.62 (s, 3H), 1.04 – 0.78 (m, 5H), 0.60 – 0.50 (m, 1H). **^13^C NMR** (151 MHz, DMSO-*d*_6_) δ 172.5, 169.8, 163.0, 155.1, 149.8, 137.9, 136.8, 135.2, 132.3, 131.3, 130.7, 130.2, 129.9, 129.6, 128.5, 69.0, 68.9, 53.9, 38.7, 38.6, 37.5, 37.4, 33.2, 31.3, 28.0, 27.0, 24.1, 24.1, 23.0, 22.9, 14.0, 12.7, 11.3. **HRMS**: (ESI, m/z): calcd. [M+H]^+^ for C_33_H_40_ClN_6_O_3_S^+^: 635.25656, found: 635.25757. trans:cis >20:1 by ^1^H NMR.

**Compound 3. (1R,8S,9r,E)-N-(2-(2-(2-(2-((S)-4-(4-chlorophenyl)-2,3,9-trimethyl-6H-thieno[3,2-f][1,2,4]triazolo[4,3-a][1,4]diazepin-6-yl)acetamido)ethoxy)ethoxy)ethyl)bicyclo[6.1.0]non-4-ene-9-carboxamide**

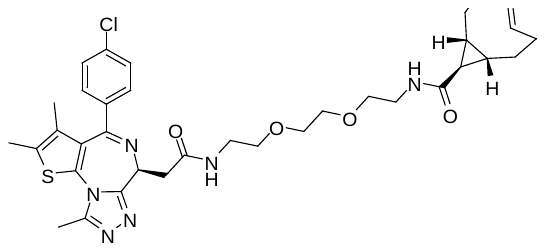

General procedure A was followed using (S)-N-(2-(2-(2-aminoethoxy)ethoxy)ethyl)-2-(4-(4-chlorophenyl)-2,3,9-trimethyl-6H-thieno[3,2-f][1,2,4]triazolo[4,3-a][1,4]diazepin-6-yl)acetamide (76 mg, 0.14 mmol) as the amine coupling partner. The crude residue was purified with a Waters HPLC system (mass triggered collection) using a Phenomenex Gemini NX-C18 225x30 mm column under NH_4_OH conditions (40-70% CH_3_CN/H_2_O with 0.1% NH_4_OH, 14 minute run, 50 mL/min), to afford the title compound **3** (27 mg, 33 % yield) as a yellow oil. **^1^H NMR** (500 MHz, DMSO-*d*_6_) δ 8.27 (t, *J* = 5.6 Hz, 1H), 7.92 (t, *J* = 5.6 Hz, 1H), 7.48 (d, *J* = 8.8 Hz, 2H), 7.42 (d J = 8.6 Hz, 2H), 5.78 (ddd, *J* = 16.7, 9.2, 6.2 Hz, 1H), 5.09 (ddd, *J* = 16.8, 10.5, 3.8 Hz, 1H), 4.50 (dd, *J* = 8.0, 6.1 Hz, 1H), 3.57 – 3.50 (m, 4H), 3.46 (t, *J* = 5.9 Hz, 2H), 3.40 (t, *J* = 5.9 Hz, 2H), 3.35 – 3.14 (m, 6H), 2.59 (s, 3H), 2.41 (s, 3H), 2.30 – 2.07 (m, 4H), 1.92 – 1.80 (m, 2H), 1.62 (s, 3H), 1.05 – 0.81 (m, 4H), 0.63 – 0.52 (m, 1H). **^13^C NMR** (151 MHz, DMSO-*d*_6_) δ 172.5, 169.7, 163.0, 155.1, 149.8, 137.9, 136.8, 135.2, 132.3, 131.4, 130.7, 130.2, 129.9, 129.6, 128.5, 69.6, 69.5, 69.2, 53.8, 48.6, 38.7, 38.7, 37.5, 37.5f, 33.2, 31.4, 28.0, 27.0, 24.1, 23.0, 14.1, 12.7, 11.3. **HRMS**: (ESI, m/z): calcd. [M+H]^+^ for C_35_H_44_ClN_6_O_4_S^+^: 679.28278, found: 679.28369. trans:cis >20:1 by ^1^H NMR.

**Compound 4. (1R,8S,9r,E)-N-(1-((S)-4-(4-chlorophenyl)-2,3,9-trimethyl-6H-thieno[3,2-f][1,2,4]triazolo[4,3-a][1,4]diazepin-6-yl)-2-oxo-6,9,12-trioxa-3-azatetradecan-14-yl)bicyclo[6.1.0]non-4-ene-9-carboxamide**

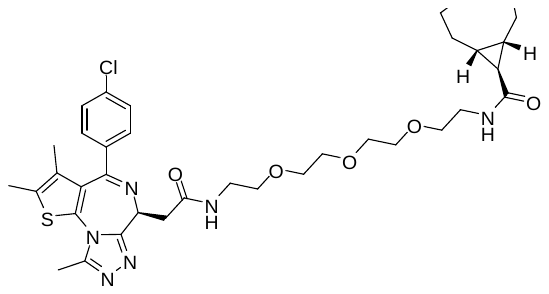

General procedure A was followed using (S)-N-(2-(2-(2-(2-aminoethoxy)ethoxy)ethoxy)ethyl)-2-(4-(4-chlorophenyl)-2,3,9-trimethyl-6H-thieno[3,2-f][1,2,4]triazolo[4,3-a][1,4]diazepin-6-yl)acetamide (83 mg, 0.14 mmol) as the amine coupling partner. The crude residue was purified with a Waters HPLC system (mass triggered collection) using a Phenomenex Gemini NX-C18 225x30 mm column under NH_4_OH conditions (40-70% CH_3_CN/H_2_O with 0.1% NH_4_OH, 14 minute run, 50 mL/min), to afford the title compound **4** (53 mg, 61 % yield) as a yellow oil. **^1^H NMR** (500 MHz, DMSO-*d*_6_): δ 8.26 (t, *J* = 5.7 Hz, 1H), 7.92 (t, *J* = 5.6 Hz, 1H), 7.48 (d, *J* = 8.7 Hz, 2H), 7.42 (d, *J* = 8.6 Hz, 2H), 5.79 (ddd, *J* = 16.8, 9.2, 6.2 Hz, 1H), 5.10 (ddd, *J* = 16.9, 10.5, 3.8 Hz, 1H), 4.50 (dd, J = 8.1, 6.0 Hz, 1H), 3.55 – 3.48 (m, 8H), 3.45 (t, J = 5.9 Hz, 2H), 3.39 (t, J = 5.9 Hz, 2H) 3.30 – 3.14 (m, 6H), 2.59 (s, 3H), 2.41 (s, 3H), 2.28 – 2.22 (m, 1H), 2.21 – 2.09 (m, 3H), 1.93 – 1.82 (m, 2H), 1.62 (s, 3H), 1.04 – 0.95 (m, 1H), 0.93 – 0.81 (m, 3H), 0.63 – 0.52 (m, 1H). **^13^C NMR** (151 MHz, DMSO-*d*_6_) δ 172.5, 169.7, 163.0, 155.1, 149.8, 137.9, 136.8, 135.2, 132.3, 131.4, 130.7, 130.2, 129.9, 129.6, 128.5, 69.8, 69.7, 69.6, 69.5, 69.2, 69.2, 53.8, 38.8, 38.6, 37.5, 37.5, 33.2, 31.4, 28.0, 27.0, 24.1, 23.0, 14.1, 12.7, 11.3. **HRMS**: (ESI, m/z): calcd. [M+H]^+^ for C_37_H_48_ClN_6_O_5_S^+^: 723.30899, found: 723.31012. trans:cis >20:1 by ^1^H NMR.

**Compound 5. (1R,8S,9r,E)-N-(1-((S)-4-(4-chlorophenyl)-2,3,9-trimethyl-6H-thieno[3,2-f][1,2,4]triazolo[4,3-a][1,4]diazepin-6-yl)-2-oxo-6,9,12,15-tetraoxa-3-azaheptadecan-17-yl)bicyclo[6.1.0]non-4-ene-9-carboxamide**

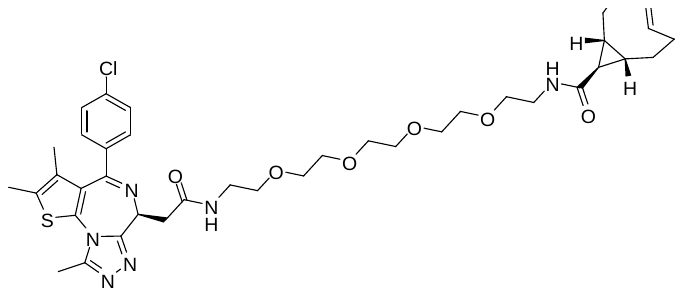

General procedure A was followed using (S)-N-(14-amino-3,6,9,12-tetraoxatetradecyl)-2-(4-(4-chlorophenyl)-2,3,9-trimethyl-6H-thieno[3,2-f][1,2,4]triazolo[4,3-a][1,4]diazepin-6-yl)acetamide (89 mg, 0.14 mmol) as the amine coupling partner. The crude residue was purified with a Waters HPLC system (mass triggered collection) using a Phenomenex Gemini NX-C18 225x30 mm column under NH_4_OH conditions (40-70% CH_3_CN/H_2_O with 0.1% NH_4_OH, 14 minute run, 50 mL/min), to afford the title compound **5** (20 mg, 21 % yield) as a yellow oil. **^1^H NMR** (500 MHz, DMSO-*d*_6_) δ 8.26 (t, *J* = 5.7 Hz, 1H), 7.92 (t, *J* = 5.6 Hz, 1H), 7.48 (d, *J* = 8.8 Hz, 2H), 7.42 (d, *J* = 8.6 Hz, 2H), 5.79 (ddd, *J* = 16.1, 9.3, 6.2 Hz, 1H), 5.10 (ddd, *J* = 16.9, 10.4, 3.8 Hz, 1H), 4.50 (dd, *J* = 8.1, 6.0 Hz, 1H), 3.56 – 3.43 (m, 14H), 3.38 (t, *J* = 5.9 Hz, 2H), 3.31 – 3.13 (m, 6H), 2.59 (s, 3H), 2.41 (s, 3H), 2.29 – 2.1 (m, 4H), 1.94 – 1.80 (m, 2H), 1.62 (s, 3H), 1.04 – 0.95 (m, 1H), 0.94 – 0.91 (m, 3H), 0.63 – 0.53 (m, 1H). **^13^C NMR** (151 MHz, DMSO-*d*_6_) δ 172.4, 169.7, 163.0, 155.1, 149.8, 137.9, 136.8, 135.2, 132.3, 131.4, 130.7, 130.1, 129.8, 129.6, 128.5, 69.8, 69.8, 69.8, 69.7, 69.6, 69.5, 69.2, 69.2, 53.8, 38.7, 38.6, 37.5, 37.5, 33.2, 31.4, 28.0, 27.0, 24.1, 22.9, 14.0, 12.7, 11.3. **HRMS**: (ESI, m/z): calcd. [M+H]^+^ for C_39_H_52_ClN_6_O_6_S^+^: 767.33521, found: 767.33594. trans:cis >20:1 by ^1^H NMR.

**Compound 6. (1R,8S,9r,E)-N-(1-((S)-4-(4-chlorophenyl)-2,3,9-trimethyl-6H-thieno[3,2-f][1,2,4]triazolo[4,3-a][1,4]diazepin-6-yl)-2-oxo-6,9,12,15,18-pentaoxa-3-azaicosan-20-yl)bicyclo[6.1.0]non-4-ene-9-carboxamide**

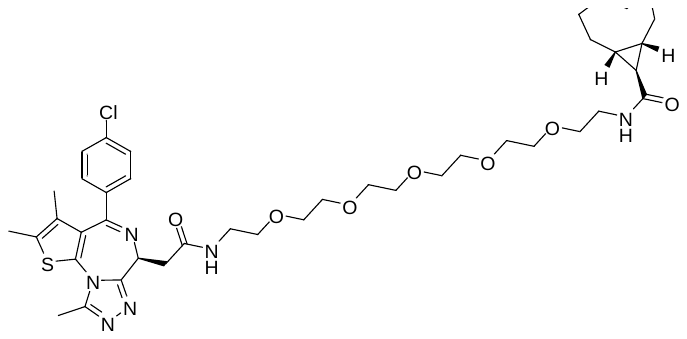

General procedure A was followed using (S)-N-(17-amino-3,6,9,12,15-pentaoxaheptadecyl)-2-(4-(4-chlorophenyl)-2,3,9-trimethyl-6H-thieno[3,2-f][1,2,4]triazolo[4,3-a][1,4]diazepin-6-yl)acetamide (96 mg, 0.14 mmol) as the amine coupling partner. The crude residue was purified with a Waters HPLC system (mass triggered collection) using a Phenomenex Gemini NX-C18 225x30 mm column under ammonium hydroxide conditions (45-75% CH_3_CN/H_2_O with 0.1% NH_4_OH, 14 minute run, 50 mL/min), to afford the title compound **6** (32 mg, 32 % yield) as a yellow oil. **^1^H NMR** (400 MHz, DMSO-*d*_6_) δ 8.26 (t, *J* = 5.6 Hz, 1H), 7.91 (t, *J* = 5.6 Hz, 1H), 7.48 (d, *J* = 8.7 Hz, 2H), 7.42 (d, *J* = 8.6 Hz, 2H), 5.79 (ddd, *J* = 16.1, 9.2, 6.2 Hz, 1H), 5.11 (ddd, *J* = 17.0, 10.4, 3.8 Hz, 1H), 4.50 (dd, *J* = 8.1, 6.0 Hz, 1H), 3.56 – 3.42 (m, 19H), 3.38 (t, *J* = 5.9 Hz, 2H), 2.59 (s, 3H), 2.41 (s, 3H), 2.31 – 2.09 (m, 4H), 1.95 – 1.79 (m, 2H), 1.62 (s, 3H), 1.04 – 0.80 (m, 4H), 0.65 – 0.52 (m, 1H). **^13^C NMR** (151 MHz, DMSO-*d*_6_) δ 172.4, 169.7, 163.0, 155.1, 149.8, 137.9, 136.8, 135.2, 132.3, 131.3, 130.7, 130.1, 129.8, 129.6, 128.4, 69.8, 69.8, 69.7, 69.7, 69.6, 69.5, 69.2, 69.2, 58.8, 38.8, 38.6, 37.5, 37.4, 33.2, 31.4, 28.0, 27.0, 24.1, 22.9, 14.0, 12.7, 11.3. **HRMS**: (ESI, m/z): calcd. [M+H]^+^ for C_41_H_56_ClN_6_O_7_S^+^: 811.36142, found: 811.36255. trans:cis >20:1 by ^1^H NMR.

**Compound 7. (1R,8S,9r,E)-N-(2-(2-((R)-4-(4-chlorophenyl)-2,3,9-trimethyl-6H-thieno[3,2-f][1,2,4]triazolo[4,3-a][1,4]diazepin-6-yl)acetamido)ethyl)bicyclo[6.1.0]non-4-ene-9-carboxamide**

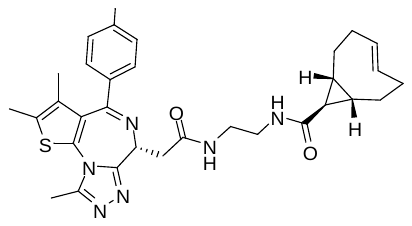

General Procedure A was followed using 0.06 mmol (10 mg) of sTCO acid and (R)-N^1^-(2-(4-(4-chlorophenyl)-2,3,9-trimethyl-6H-thieno[3,2-f][1,2,4]triazolo[4,3-a][1,4]diazepin-6-yl)ethyl)ethane-1,2-diamine (32 mg, 0.07 mmol) as the amine coupling partner. The crude residue was purified with a Waters HPLC system (mass triggered collection) using a Phenomenex Gemini NX-C18 225x30 mm column under NH_4_OH conditions (45-75% CH_3_CN/H_2_O with 0.1% NH_4_OH, 14 minute run, 50 mL/min), to afford the title compound **7** (7.3 mg, 21 % yield) as a yellow oil. **^1^H NMR** (500 MHz, DMSO-*d*_6_) δ 8.23 (s, 1H), 7.89 (s, 1H), 7.48 (d, *J* = 8.8 Hz, 2H), 7.42 (d, *J* = 8.6 Hz, 2H), 5.80 (ddd, *J* = 16.1, 9.2, 6.2 Hz, 1H), 5.12 (ddd, *J* = 16.9, 10.5, 3.8 Hz, 1H), 4.50 (t, *J* = 7.1 Hz, 1H), 3.23 (dd, *J* = 7.1, 2.6 Hz, 2H), 3.14 (s, 4H), 2.59 (s, 3H), 2.41 (s, 3H), 2.29 – 2.10 (m, 4H), 1.94 – 1.81 (m, 2H), 1.62 (s, 3H), 1.06 – 0.96 (m, 1H), 0.95 – 0.81 (m, 3H), 0.66 – 0.53 (m, 1H),. **^13^C NMR** (151 MHz, DMSO-*d*_6_) δ 172.6, 169.8, 163.1, 155.1, 149.9, 137.9, 136.8, 135.2, 132.3, 131.4, 130.7, 130.2, 129.9, 129.6, 128.5, 53.8, 38.7, 38.6, 37.6, 37.5, 33.2, 31.4, 28.2, 27.0, 24.2, 24.2, 23.1, 23.0, 14.1, 12.7, 11.3. **HRMS**: (ESI, m/z): calcd. [M+H]^+^ for C_31_H_36_ClN_6_O_2_S^+^: 591.23035, found: 591.23138. trans:cis >20:1 by ^1^H NMR.

**Compound 8. (1R,8S,9r,E)-N-(2-(2-(2-((R)-4-(4-chlorophenyl)-2,3,9-trimethyl-6H-thieno[3,2-f][1,2,4]triazolo[4,3-a][1,4]diazepin-6-yl)acetamido)ethoxy)ethyl)bicyclo[6.1.0]non-4-ene-9-carboxamide**

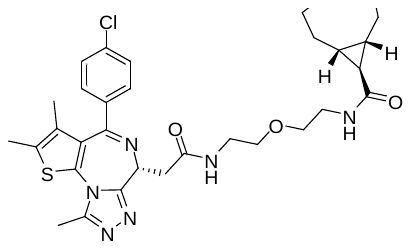

General Procedure A was followed using (R)-N-(2-(2-aminoethoxy)ethyl)-2-(4-(4-chlorophenyl)-2,3,9-trimethyl-6H-thieno[3,2-f][1,2,4]triazolo[4,3-a][1,4]diazepin-6-yl)acetamide (68 mg, 0.14 mmol) as the amine coupling partner. The crude residue was purified with a Waters HPLC system (mass triggered collection) using a Phenomenex Gemini NX-C18 225x30 mm column under NH_4_OH conditions (40-70% CH_3_CN/H_2_O with 0.1% NH_4_OH, 14 minute run, 50 mL/min), to afford the title compound **8** (46 mg, 60 % yield) **^1^H NMR** (400 MHz, DMSO-*d*_6_) δ 8.26 (t, *J* = 5.6 Hz, 1H), 7.92 (t, *J* = 5.5 Hz, 1H), 7.48 (d, *J* = 8.7 Hz, 2H), 7.42 (d, *J* = 8.6 Hz, 2H), 5.84 – 5.71 (m, 1H), 5.08 (ddd, *J* = 15.7, 10.3, 3.9 Hz, 1H), 4.50 (dd, *J* = 8.3, 5.9 Hz, 1H), 3.47 – 3.15 (m, 10H presumed under solvent peak), 2.59 (s, 3H), 2.41 (s, 3H), 2.29 – 2.05 (m, 4H), 1.91 – 1.76 (m, 2H), 1.62 (s, 3H), 1.05 – 0.76 (m, 4H), 0.55 (q, *J* = 12.5 Hz, 1H). **^13^C NMR** (151 MHz, DMSO-*d*_6_) δ 172.5, 169.8, 163.1, 155.1, 149.8, 137.9, 136.8, 135.2, 132.3, 131.4, 130.7, 130.2, 129.9, 129.6, 128.5, 69.0, 68.9, 53.9, 38.7, 38.6, 37.5, 37.4, 33.2, 31.3, 28.1, 27.0, 24.1, 24.1, 23.0, 23.0, 14.1, 12.7, 11.3. **HRMS**: (ESI, m/z): calcd. [M+H]^+^ for C_33_H_40_ClN_6_O_3_S^+^: 635.25656, found: 635.25769. trans:cis >20:1 by ^1^H NMR.

1.	Marjanovic J, Baranczak A, Marin V, Stockmann H, Richardson PL, Vasudevan A. Development of inverse electron demand Diels-Alder ligation and TR-FRET assays for the determination of ligand-protein target occupancy in live cells. *MedChemComm* **8**, 789-795 (2017).

2.	Herner A*, et al.* 2-Aryl-5-carboxytetrazole as a New Photoaffinity Label for Drug Target Identification. *J Am Chem Soc* **138**, 14609-14615 (2016).

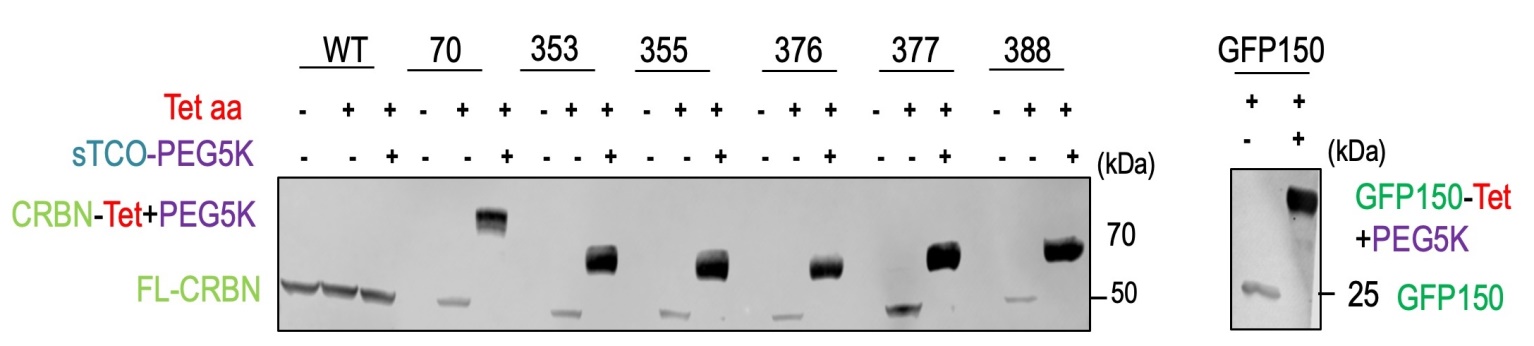

**Fig. S1:** **Expression and ligation of CRBN^Tet^ forms** **in CRBN K/O cells.** (a) Similar to Fig. 3b, shown is an annotated FLAG-tag-based immunoblot for CRBN variants (left hand gel) and sfGFP150 control (right hand gel) documenting expression levels in HEK293T CRBN K/O cells, and also showing reactivity based on mobility shift after *in vitro* reaction with sTCO-PEG_5K_. GAPDH loading level control included. Cells were transfected with 600 ng of the respective CRBN-expressing plasmid DNA and grown with 30 µM Tet3Bu for at least 36 h.

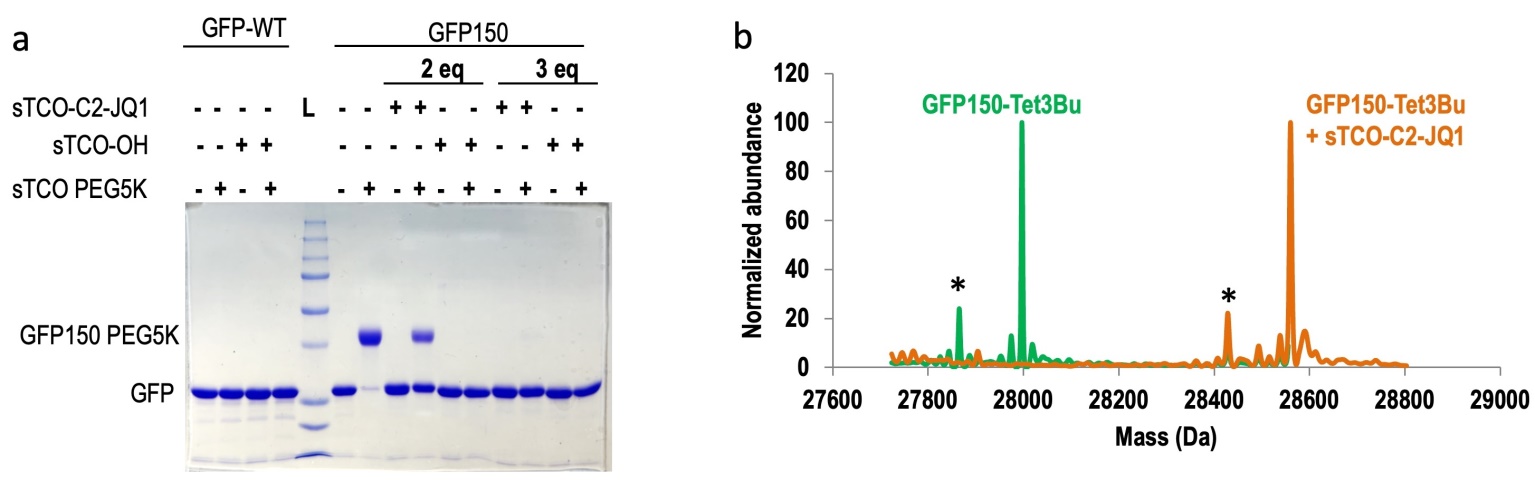

**Fig. S2: *In vitro* reactivity of purified Tet3Bu protein with sTCO-target ligand.** His-tagged GFP^WT^ and Tet3Bu-containing sfGFP^150^ protein were purified from BL21 (DE3) cells grown in autoinduction media co transformed with the Mb-pAJE-Tet3.0 and pET28-sfGFP-150(TAG) or pET28-GFP^WT^ plasmids. The sfGFP^150^ was produced from cells grown in 0.5 mM Tet3Bu. (a) Mobility shift assay gel with lane 5 having molecular weight standards and the conditions of the others lanes as annotated. For lanes 2, 4 and 7 the GFP form was incubated with 10 eq. of sTCO-PEG_5K_ for 5 min prior to adding SDS load buffer, and noteworthy is the ~90 % mobility shift in lane 7 of the Tet-sfGFP^150^ protein and not GFP^WT^, which confirms the reactivity of encoded Tet3Bu. For lanes 8-11, the purified Tet-sfGFP^150^ was incubated with 2 eq. of sTCO-C2-JQ1 and sTCO-OH for 5 min. As seen in lane 9, subsequent addition of sTCO-PEG_5K_ showed a shift for ~50% of the protein meaning that the sTCO-C2-JQ1 left ~50% of the protein unmodified. Lanes 12-15 show a treatment with 3 eq. of sTCO-C2-JQ1 or sTCO-OH r for 15 min prior to addition of sTCO-PEG_5K_ and the lack of a shift seen in lane 13 means that this led to complete reactivity of sTCO-JQ1 degrader ligand with the Tet-GFP^150^. **(b)** Mass spectrometry results for the purified Tet-sfGFP^150^ protein (green) and the protein after reaction sTCO-C2-JQ1 (orange). Each sample has the appropriate mass for the expected product (Tet-sfGFP^150^ predicted: 27996.25 Da avg, observed: 27996.4 Da avg; and Tet-sfGFP^150^+sTCO-C2-JQ1 predicted: 28559.35 Da avg observed 28559.3 Da avg. The lower mass peaks in each spectrum labeled with an asterisk match the mass expected for the loss of an N-terminal Met residue.

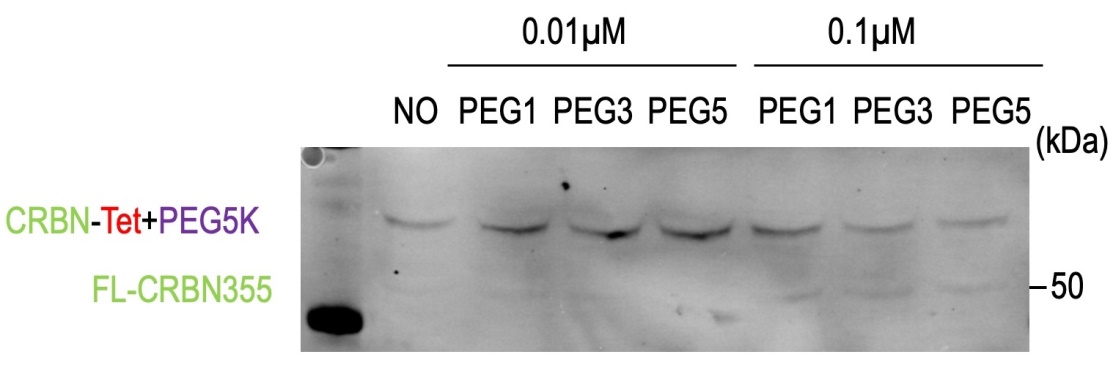

**Fig S3** **In-cell ligation of CRBN^355^ is not complete at low degrader concentrations.** Annotated immunoblot, as in Panel 3d, of mobility shift assay to detect residual reactivity of CRBN^355^ after *in vivo* reaction for 6 h with either 0.01 or 0.1 µM of three representative sTCO-JQ1 degraders (PEG1, PEG3 and PEG5 linkers). The results show that minimal in-cell labeling occurs at 0.01 µM and at or 0.1 µM the in-cell labeling is still <50%

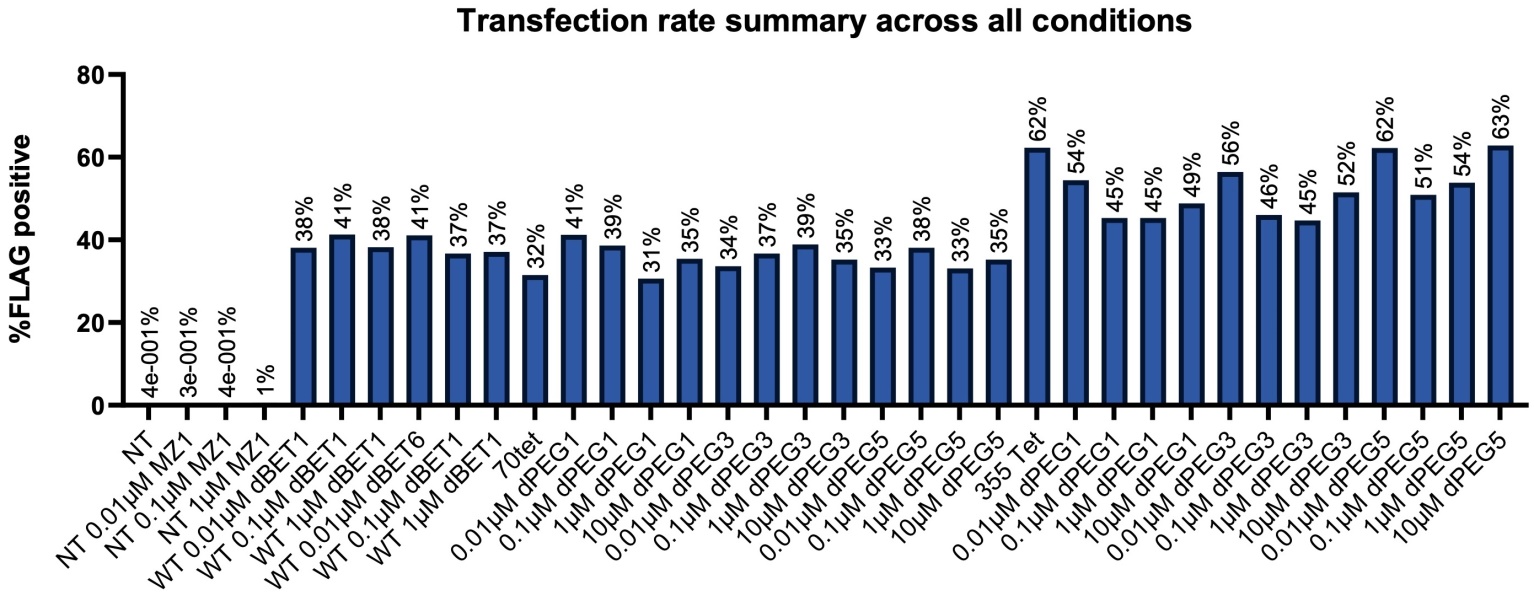

**Fig S4 Representative transfection efficiencies for cells expressing FLAG-tagged CRBN forms.** Shown is the percent of CRBN-positive cells (based on signal from the anti-FLAG antibody) for a series of representative transfections and treatment conditions as indicated on the x-axis. Transfection conditions include no transfection or transfection with the plasmid expressing either CRBN^WT^, CRBN^70^ , or CRBN^355^; treatment conditions for CRBN^WT^ include three concentrations of each dBET1 and dBET6, and for CRBN^70^ and CRBN^355^ they include four concentrations of three representative sTCO-JQ1 degraders (PEG1, PEG3 and PEG5 linkers).

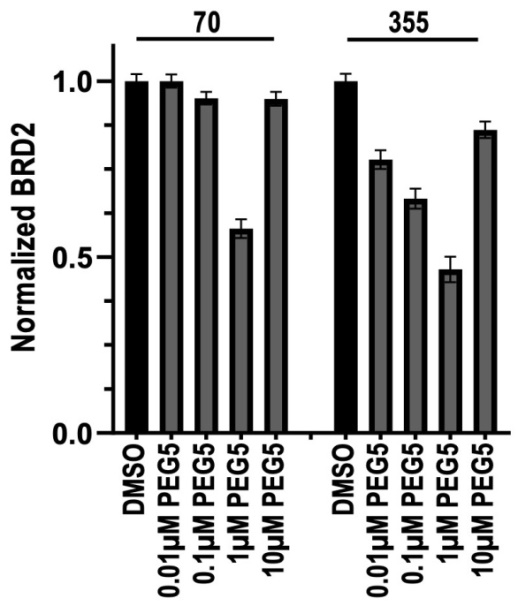

**Fig S5 Additional evidence of a higher concentration hook effect.** The fraction of remaining BRD2 seen using the IF assay is shown as a function of treatment, noted on the x-axis. Results are shown for CRBN^70^ and CRBN^355^ using the longest PEG5 ELF degrader at 0.01, 0.1, 1 and 10 µM. For both sites, the data reveal a similarly strong hook effect with much less degradation achieved at 10 µM degrader.

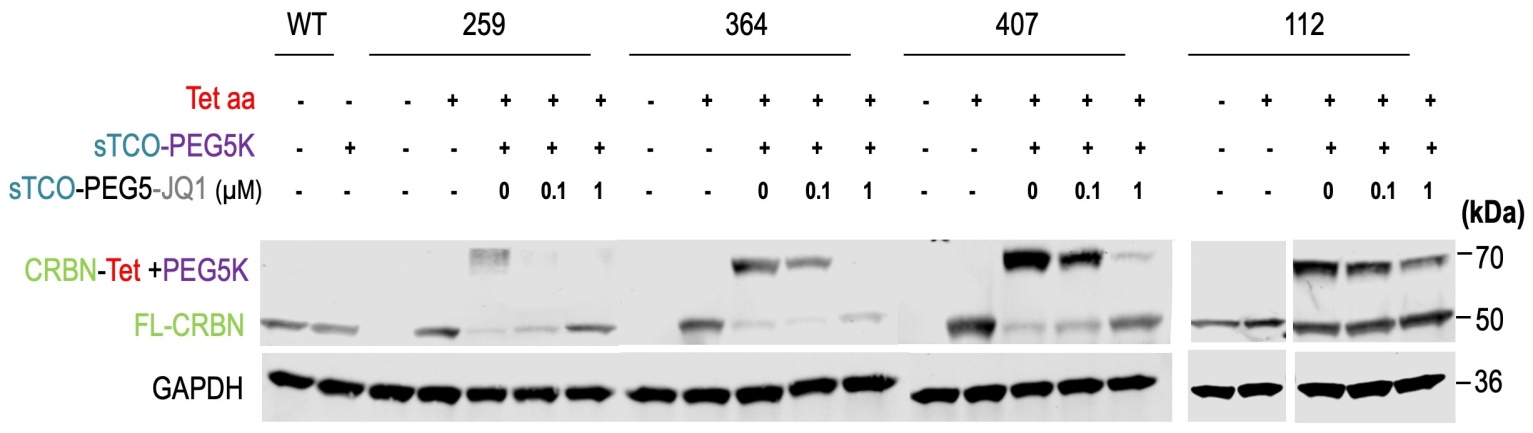

**Fig. S6: Expression and ligation of four additional CRBN^Tet^ forms in mammalian cells**. Annotated FLAG-tag-based immunoblot for CRBN variants at positions 259, 364, 407, and 112, documenting expression levels in HEK293T cells and mobility shift assay to detect residual reactivity of each CRBN^Tet^ after *in vivo* reaction for 6 h with 0.1 or 1 µM of the PEG5 sTCO-JQ1 degrader. The GAPDH loading level controls are also shown. Three of the four residues tested show good production of nearly fully reactive full-length Tet-CRBN and are fully or nearly fully labeled by sTCO degrader molecules in the cell. In contrast, CRBN^112^ expresses at reasonable levels but nearly half the produced protein does not undergo a gel-shift upon reaction with sTCO-PEG_5K_. The simplest interpretation of this is that somehow at this position the GCE system is leaky, meaning that a non-Tet amino acid is incorporated during translation. Furthermore, some of the CRBN^112^ still undergoes a gel-shift even after in-cell treatment with 1 µM degrader, meaning the in-cell labeling is not complete. Because it is only protein that has been labeled with degrader that is effective at recruiting BRD2/4 for degradation, despite the problems with the expression and the incomplete in-cell ligation, the CRBN^112^ protein should still give reliable results in this study.

**Table1 SI-1** Plasmids reported in this study.

|  | **Plasmids** | **Vector Type** | **FLAG Tag** |
| --- | --- | --- | --- |
| **1** | pAcBac1-sfGFP-WT | Mammalian pAcBac | **NO** |
| **2** | pAcBac1-sfGFP-150TAG | Mammalian pAcBac | **C-Term** |
| **3** | pAcBac1-R2-84-RS | Mammalian pAcBac | **NO** |
| **4** | pAcBac1-CRBNWT | Mammalian pAcBac | **N-Term** |
| **5** | pAcBac1-CRBN70 | Mammalian pAcBac | **N-Term** |
| **6** | pAcBac1-CRBN112 | Mammalian pAcBac | **N-Term** |
| **7** | pAcBac1-CRBN259 | Mammalian pAcBac | **N-Term** |
| **8** | pAcBac1-CRBN353 | Mammalian pAcBac | **N-Term** |
| **9** | pAcBac1-CRBN355TAG | Mammalian pAcBac | **N-Term** |
| **10** | pAcBac1-CRBN364TAG | Mammalian pAcBac | **N-Term** |
| **11** | pAcBac1-CRBN376TAG | Mammalian pAcBac | **N-Term** |
| **12** | pAcBac1-CRBN377TAG | Mammalian pAcBac | **N-Term** |
| **13** | pAcBac1-CRBN388TAG | Mammalian pAcBac | **N-Term** |
| **14** | pAcBac1-CRBN407TAG | Mammalian pAcBac | **N-Term** |

**Table SI-2**. Compounds reported in this study and their trans:cis ration as determined by ^1^H-NMR spectra.

| **Compound** | **Name** | **trans:cis** |
| --- | --- | --- |
| **1** | sTCO–C2–JQ1(+) | >20:1 |
| **2** | sTCO–PEG1–JQ1(+) | >20:1 |
| **3** | sTCO– PEG2–JQ1(+) | >20:1 |
| **4** | sTCO– PEG3–JQ1(+) | >20:1 |
| **5** | sTCO– PEG4–JQ1(+) | >20:1 |
| **6** | sTCO– PEG5–JQ1(+) | >20:1 |
| **7** | sTCO–C2–JQ1(-) | >20:1 |
| **8** | sTCO–PEG1–JQ1(-) | >20:1 |

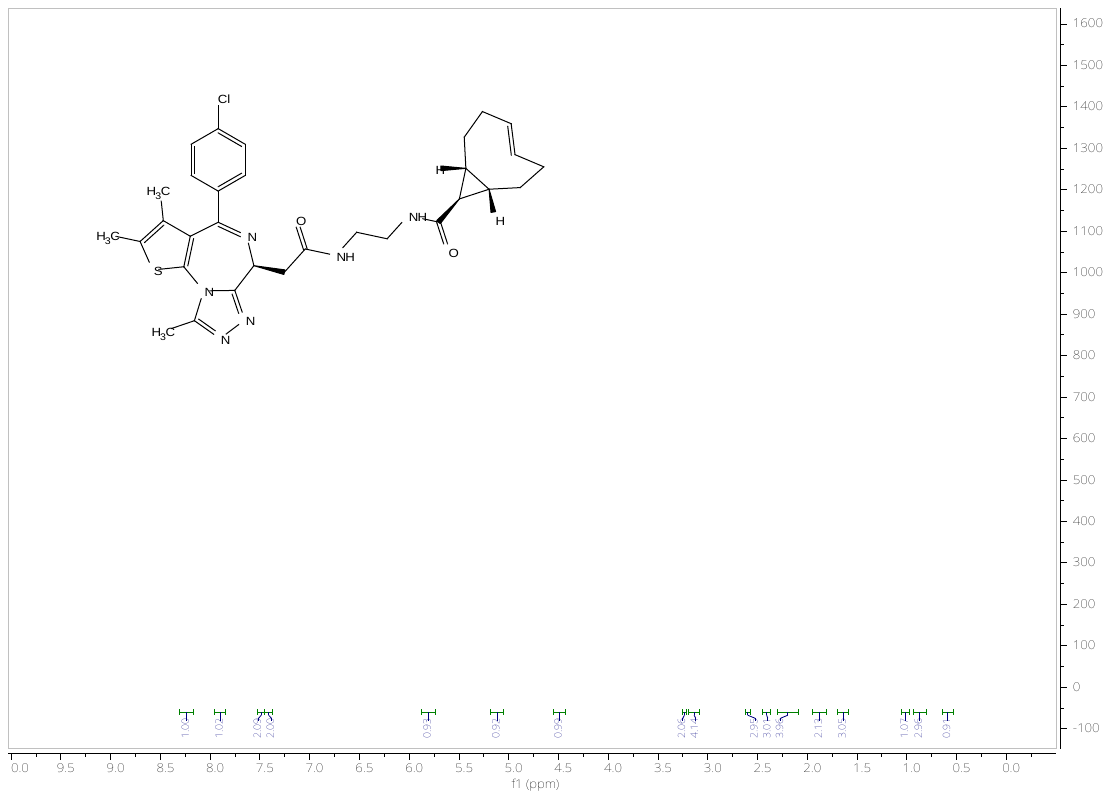
^1^H spectrum for compound **1**
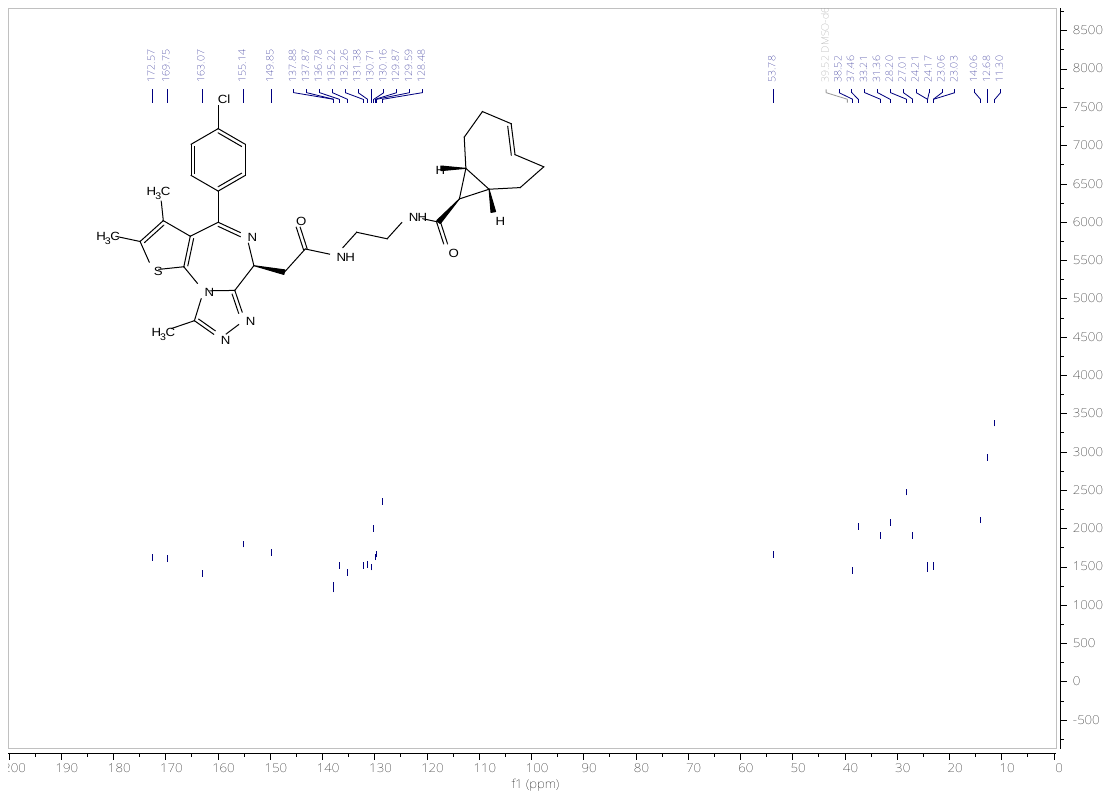

^13^C spectrum for compound **1**

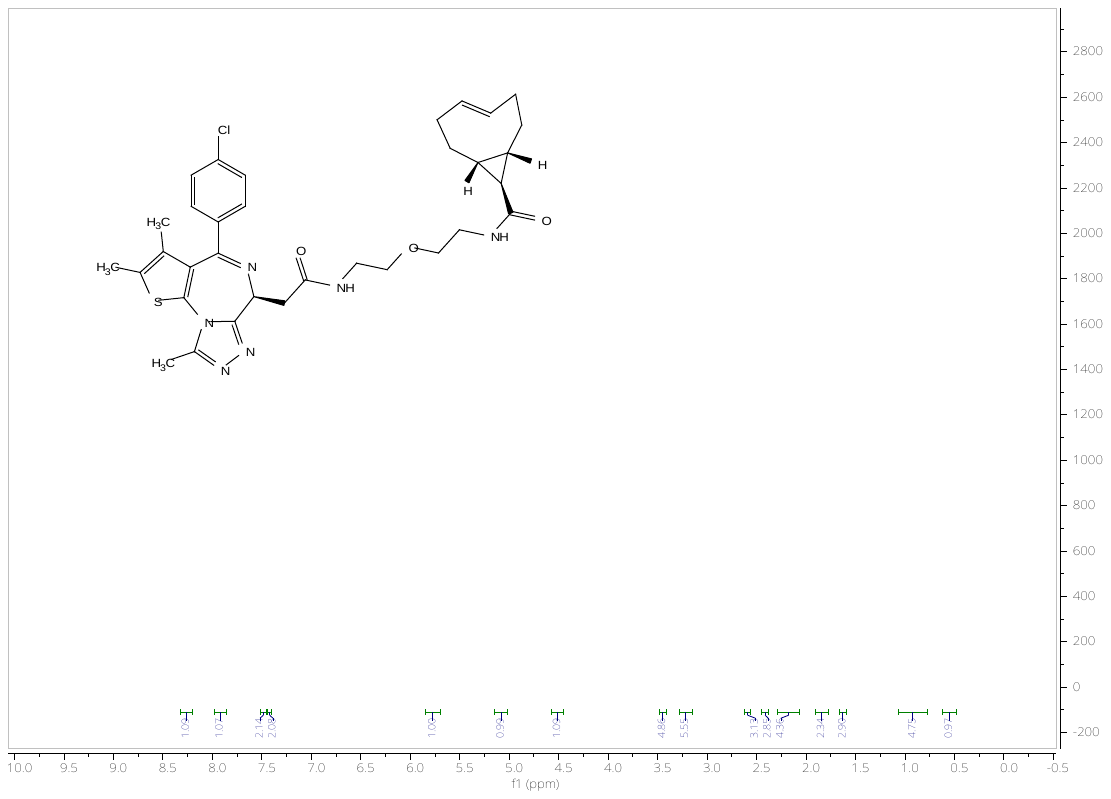
^1^H spectrum for compound **2**

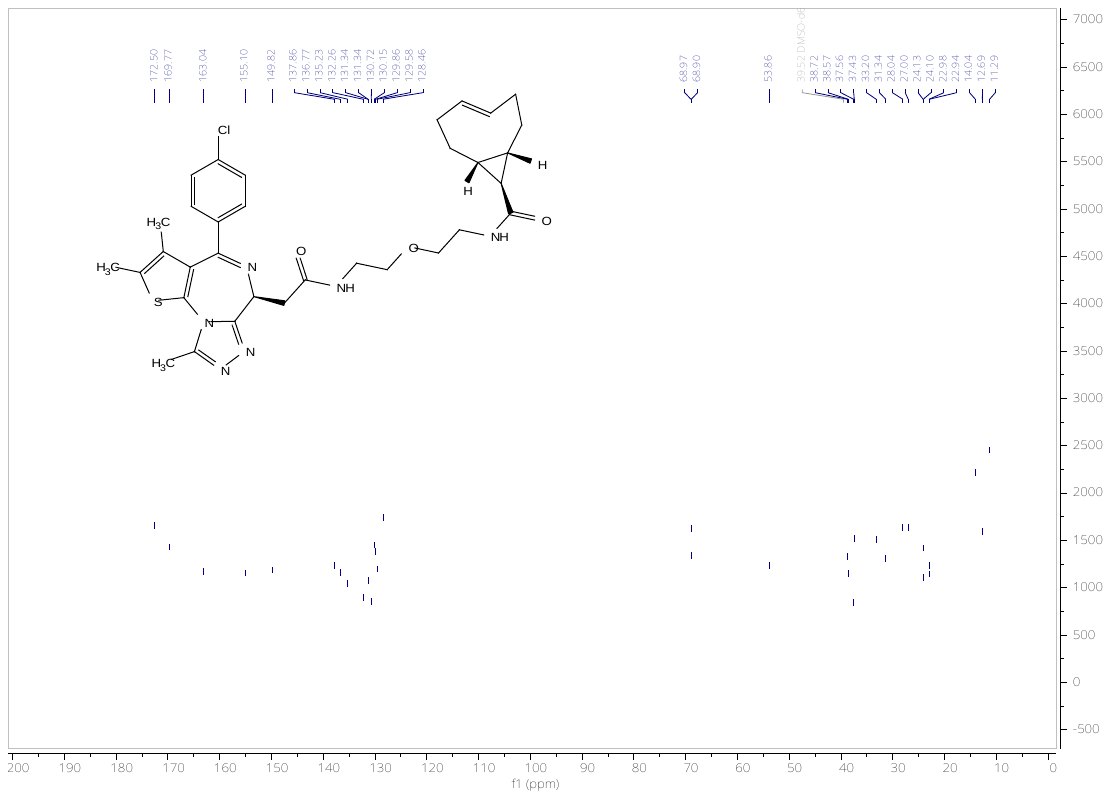
^13^C spectrum for compound **2**

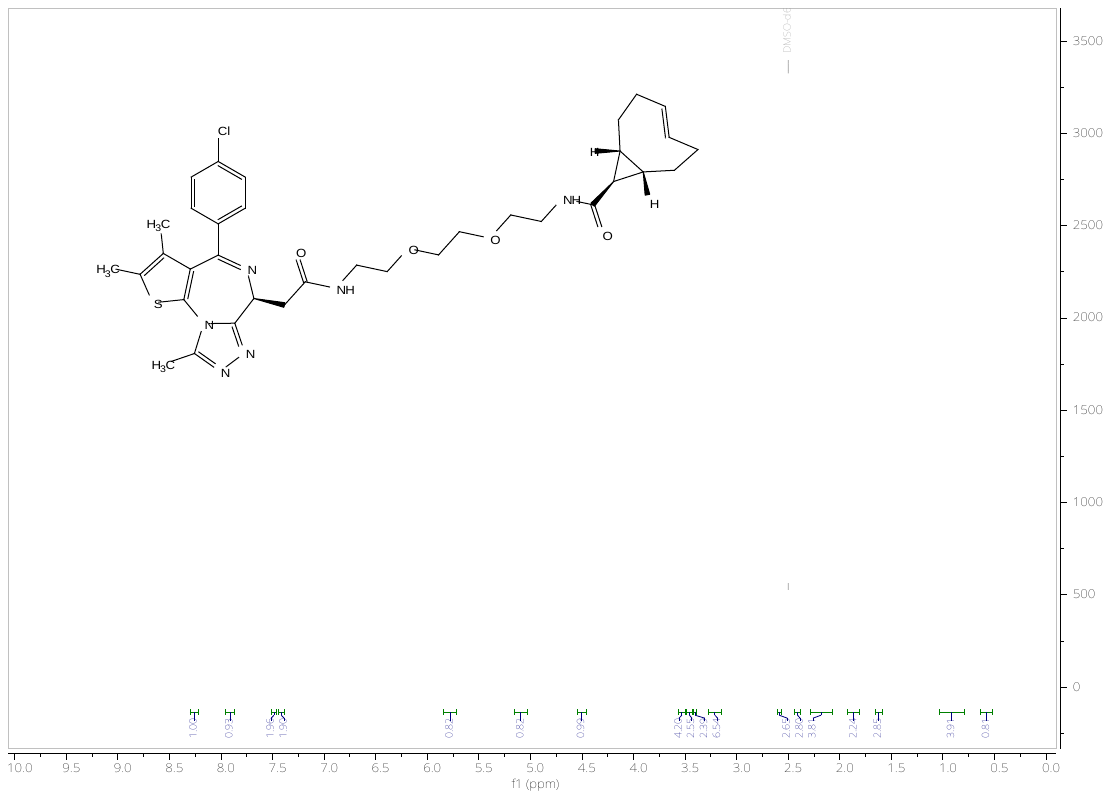
^1^H spectrum for compound **3**

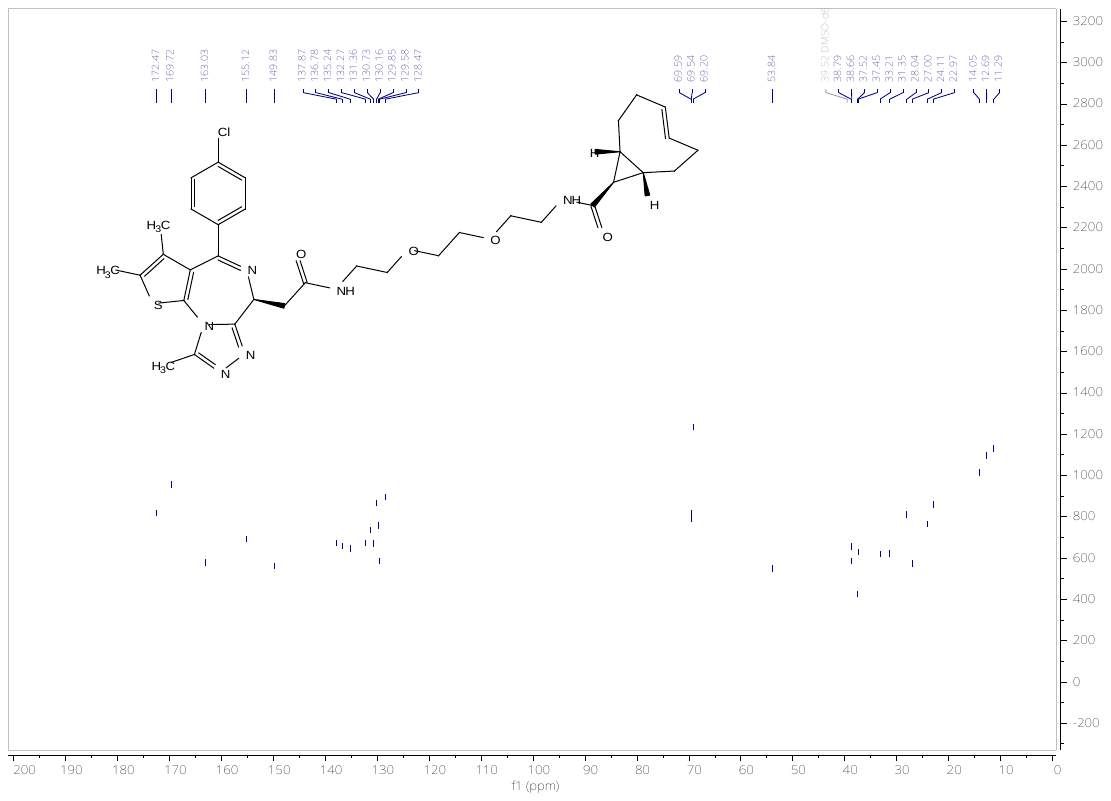

^13^C spectrum for compound **3**

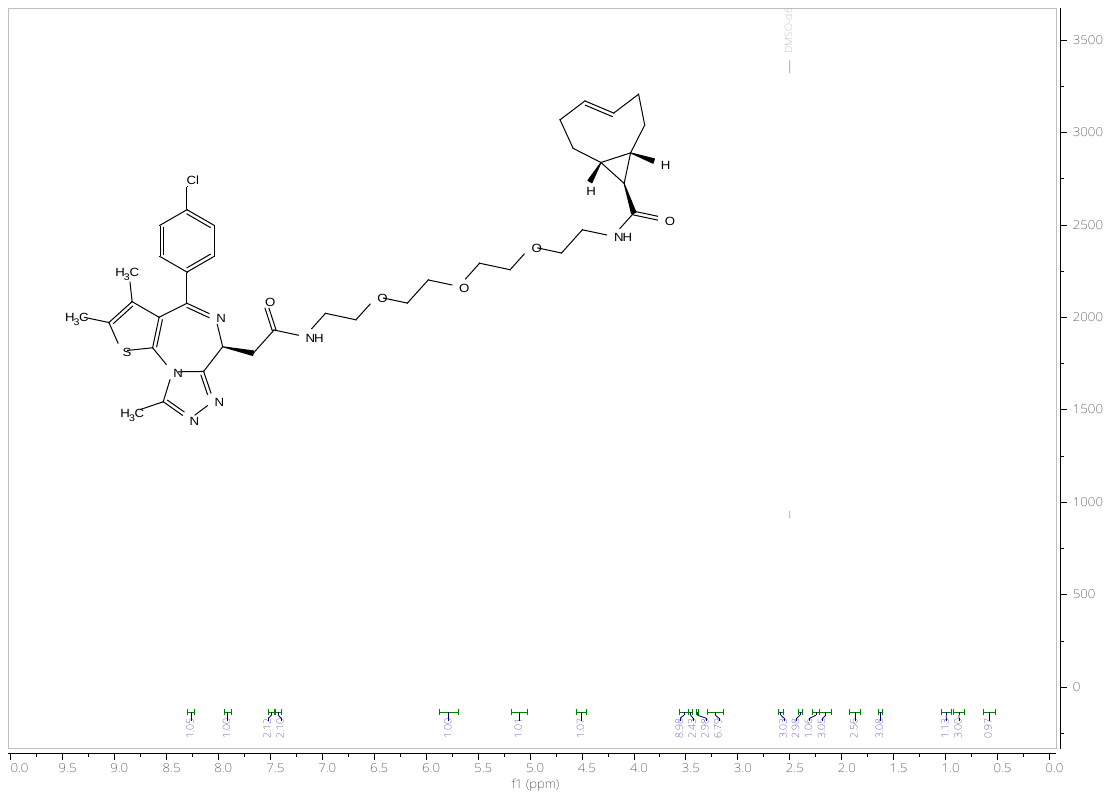
^1^H spectrum for compound **4**

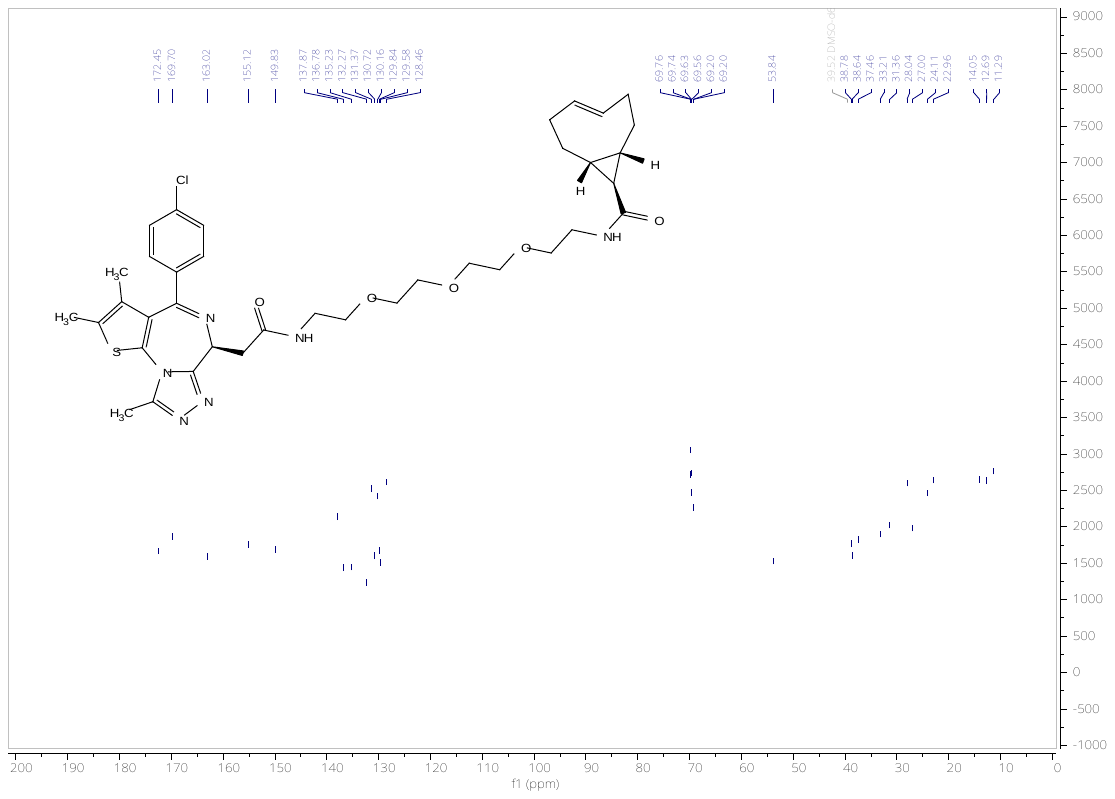
^13^C spectrum for compound **4**

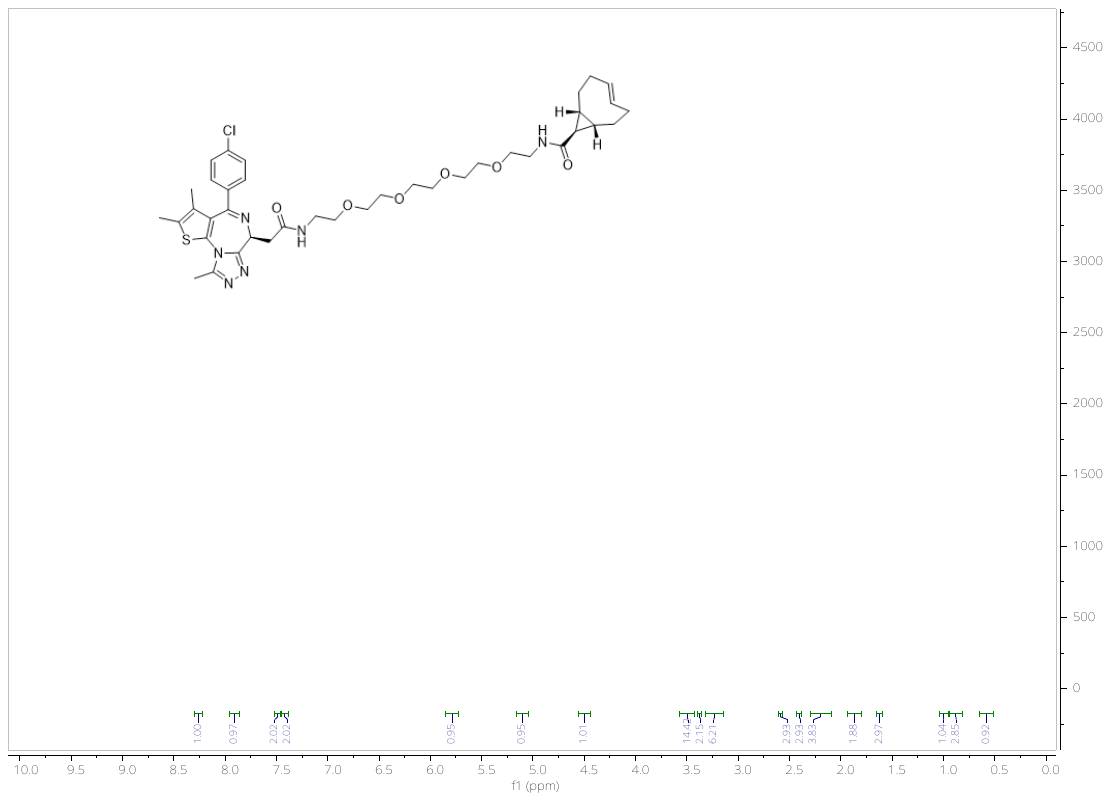
^1^H spectrum for compound **5**

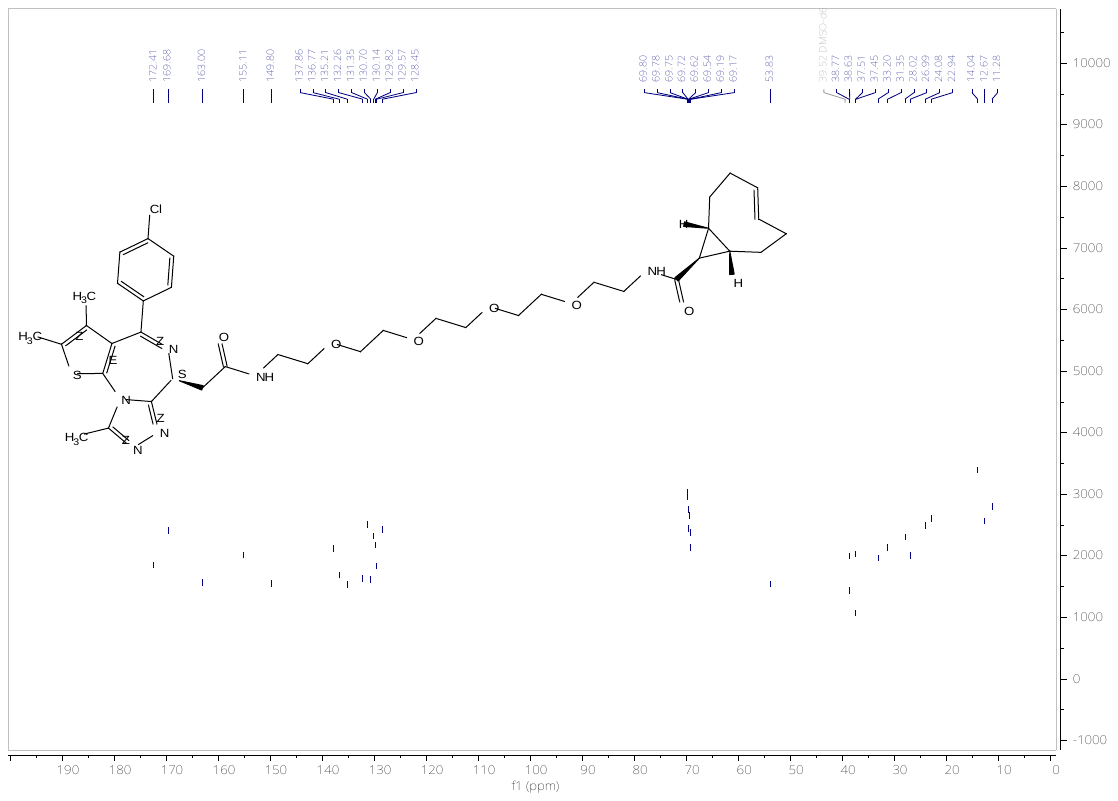

^13^C spectrum for compound **5**

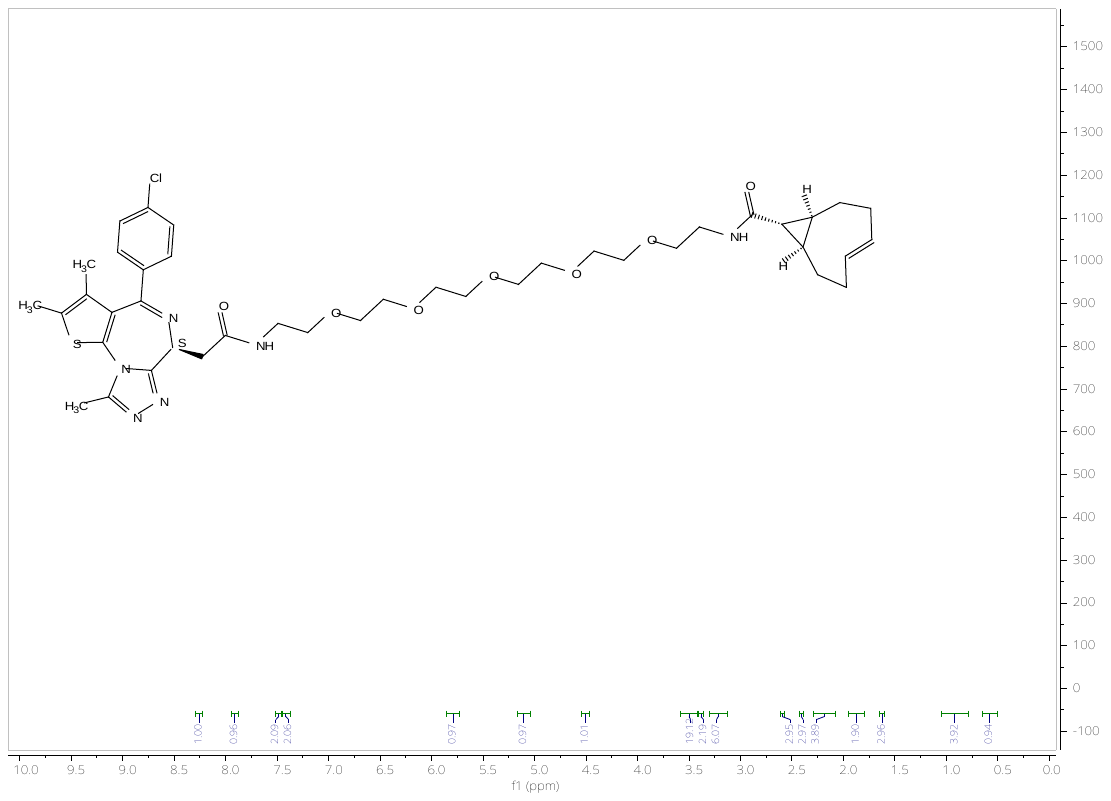
^1^H spectrum for compound **6**

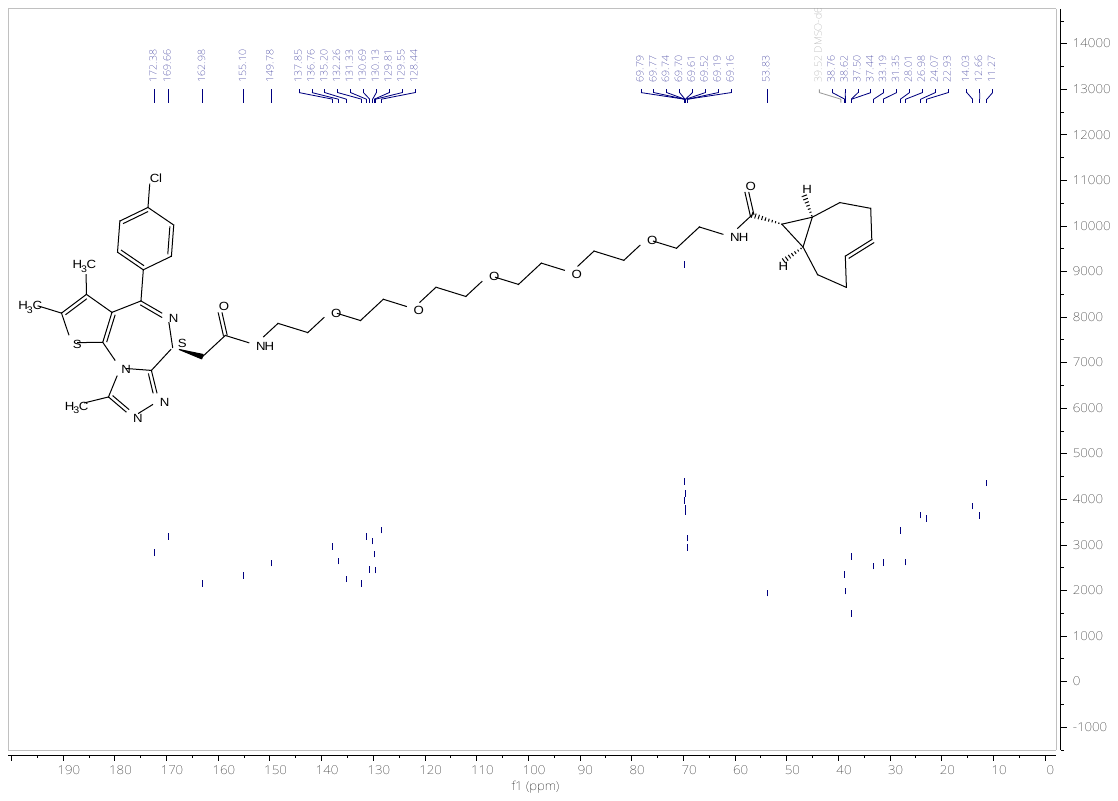
^13^C spectrum for compound **6**

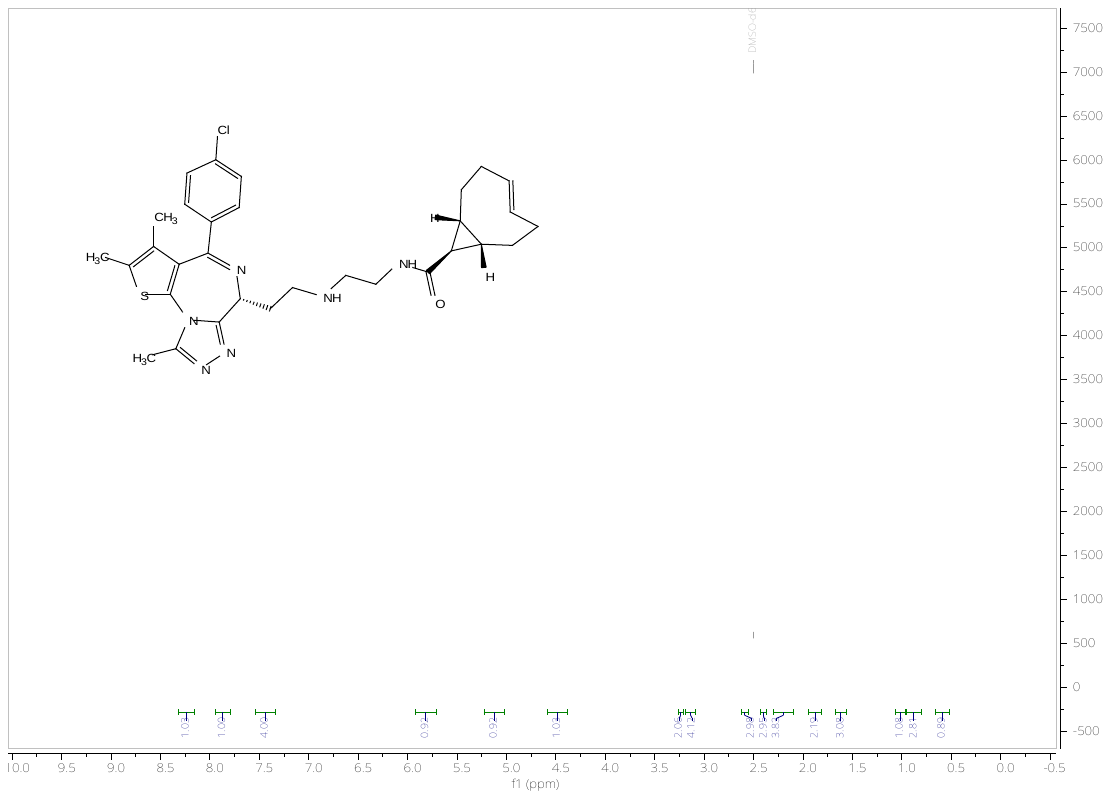
^1^H spectrum for compound **7**

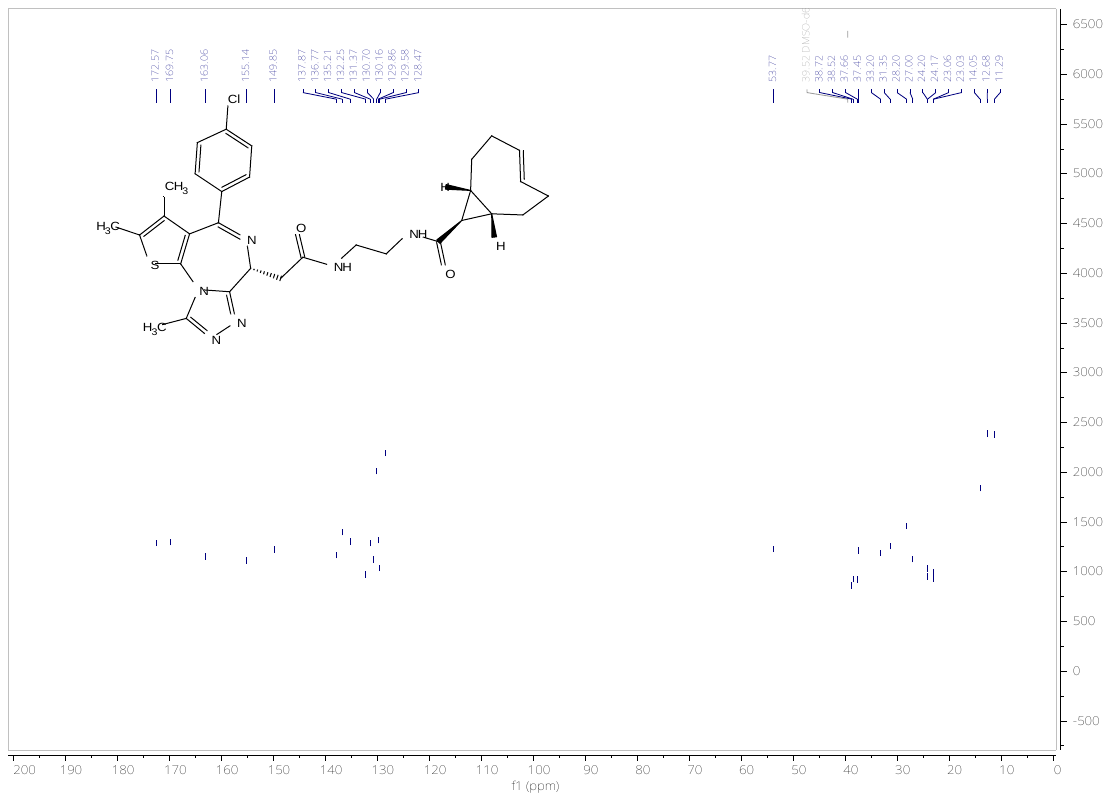
^13^C spectrum for compound **7**

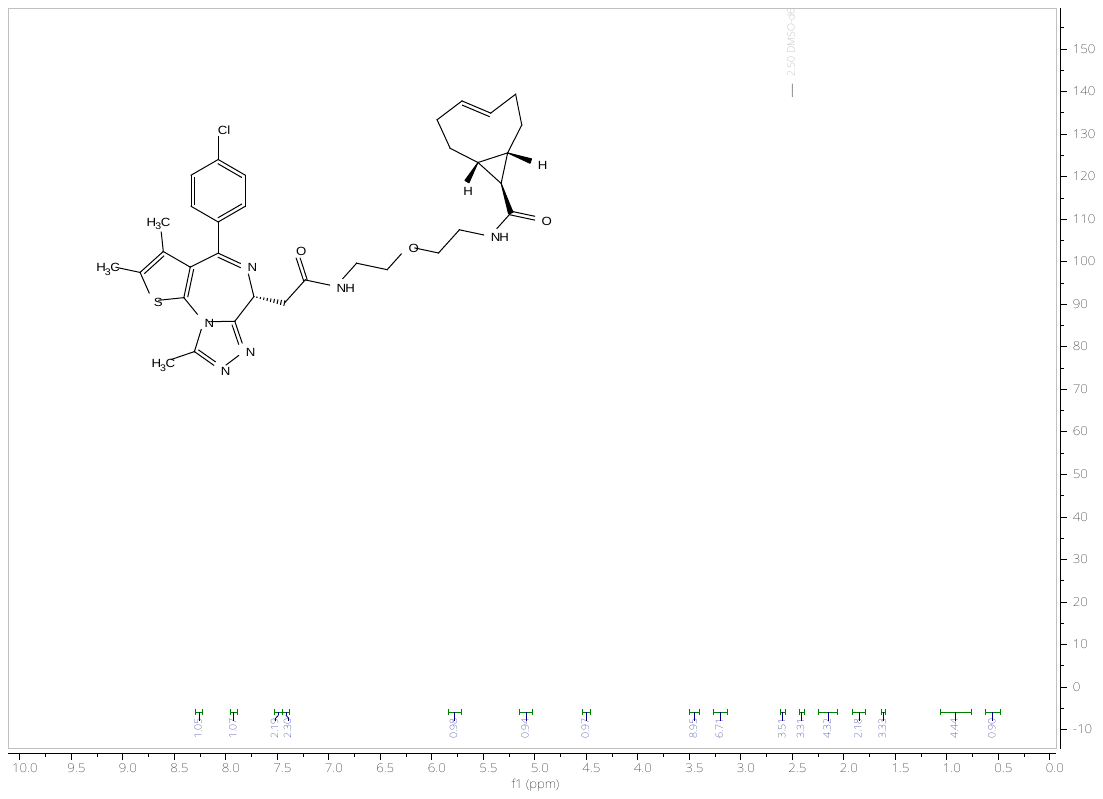
^1^H spectrum for compound **8**

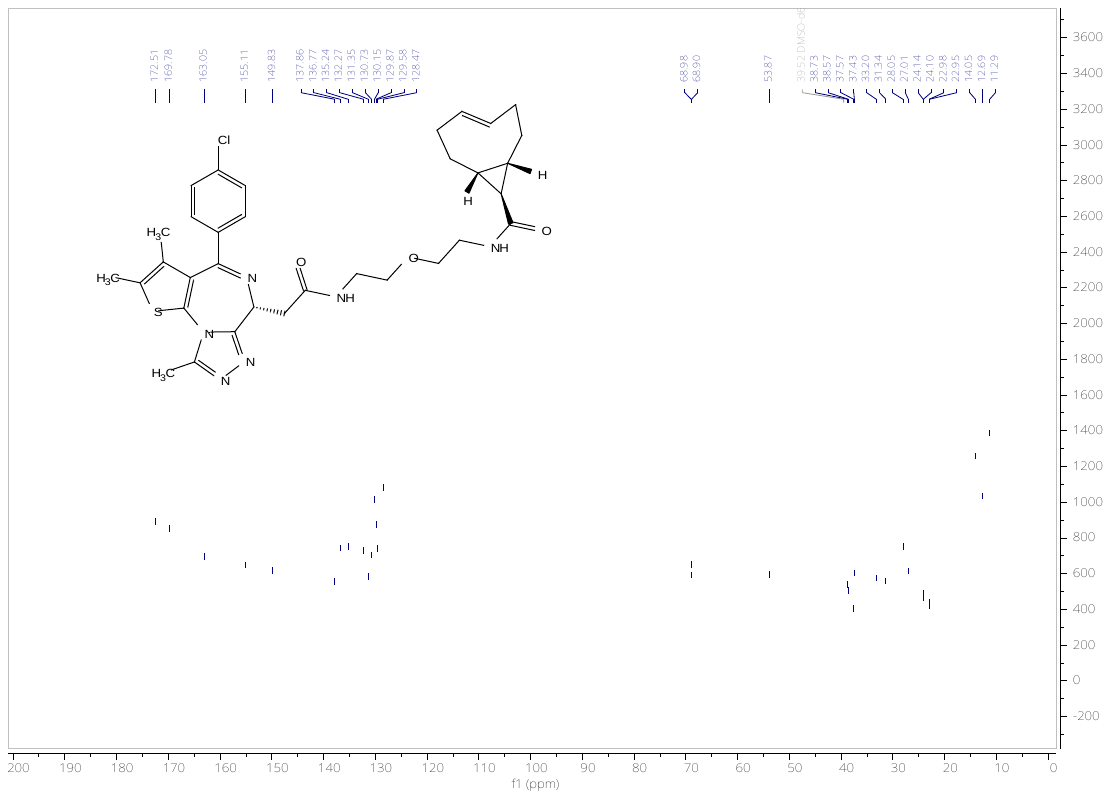

^13^C spectrum for compound **8**
